# Oral administration of dibenzoylmethane (DBM) prevents cognitive decline in a C9ORF72-mediated FTD mouse model

**DOI:** 10.64898/2026.08.07.743573

**Authors:** Paulina Torres, Daniela Becerra, José I. Astorga, Matías Fuentealba, Grant Kauwe, Luis Gonzalez, Guillermo Diaz, Vania Morales, Vicente Valenzuela, Cameron Wehrfritz, Yani Y. Ngwala, Claudia Sepulveda-Quiñenao, Samah Shah, Joanna Bons, Leonard Petrucelli, Tara E. Tracy, Birgit Schilling, Claudio Hetz

## Abstract

Amyotrophic lateral sclerosis (ALS) and frontotemporal dementia (FTD) are two related neurodegenerative disorders that display overlapping features. The hexanucleotide repeat expansion GGGGCC (G_4_C_2_) in the *C9ORF72* gene is the most common cause of ALS and FTD, which results in the accumulation of dipeptide-repeat protein aggregates. Regulation of protein synthesis at the level of the initiation factor eIF2α has been suggested as a transversal event contributing to neurodegeneration in ALS and FTD. eIF2α phosphorylation blocks protein synthesis to alleviate protein misfolding overload, but conversely it can reduce the expression of synaptic proteins resulting in neuronal dysfunction. Dibenzoylmethane (DBM) is a small molecule that reverses the translational attenuation mediated by eIF2α phosphorylation which has been shown to alleviate neurodegeneration in prion-infected mice and Tau transgenic animals. Here we investigated the efficacy of the oral administration of DBM in protecting a mouse model of C9ORF72 pathogenesis. Treatment of mice with 0.5% of DBM mixture in powdered food *ad libitum* was sufficient to prevent cognitive impairment in C9ORF72 mice. Unexpectedly, DBM treatment did not modify the content of poly(GA) and poly(GR) protein inclusion in the hippocampus and brain cortex. Proteomic profiling of brain tissue indicated that DBM administration corrected nearly 70% of the changes in gene expression triggered by expanded G_4_C_2_, where the main pathways modified by DBM were related to cytoskeleton organization, ALS, and metabolic processes. Most proteins corrected by DBM in our C9ORF72 model were also altered in the brain of human FTD/ALS patients. Overall, our results reinforce the idea that targeting protein synthesis with small molecules in patients carrying C9ORF72 mutations may result in improved cognitive capacity.

## Introduction

Amyotrophic lateral sclerosis (ALS) and frontotemporal dementia (FTD) are two related neurodegenerative diseases that are characterized by motor and cognitive impairment, respectively. ALS and FTD are part of a spectrum of diseases that overlap at the clinical, histopathological and molecular level [10, 63, 83]. ALS is a motoneuron disease characterized by progressive muscle weakness culminating in paralysis and early death [36]; however, for many years, symptoms of FTD have been reported in patients with ALS [79]. FTD patients develop personality and behavioral changes, as well as gradual impairment of language skills and cognitive function.

Protein misfolding and aggregation are key histopathological alterations of ALS/FTD [59]. In fact, the accumulation of cytoplasmic TAR DNA-binding protein 43 (TDP-43) inclusions is a central histopathological hallmark of the majority of ALS cases, as well as of FTD [14]. The hexanucleotide GGGGCC (G_4_C_2_) repeat expansion in the *C9ORF72* gene are the most frequent genetic cause of both ALS and FTD (here referred to as C9ALS/FTD) [22, 65]. C9ALS/FTD cases can harbor hundreds to thousands of G_4_C_2_ repeats, resulting on a similar probability of developing ALS or FTD [52]. This mutation leads to the expression of the repeat-associated non-ATG-mediated (RAN) peptides from different reading frames, generating “RAN” proteins of dipeptide repeats (DPR) including poly(GA), poly(GR), poly(PA), poly(GP) and poly(RP) [63, 95]. These DRP form intracellular aggregates that show different degrees of toxicity [4], highlighting poly(GA) repeats as one of the most aggregate-prone species [29].

Although multiple pathways are proposed to drive C9ALS/FTD, proteostasis disturbances arise as a significant pathogenic process [79, 80]. Relevant to this question, mRNA translation, protein synthesis, autophagy and the function of the endoplasmic reticulum (ER) are major nodes of the proteostasis network altered in C9ALS/FTD neurons [48, 67, 79]. Thus, in addition to generating abnormal protein aggregates with toxic properties, mutations in *C9ORF72* may also alter the equilibrium of the neuronal proteome, resulting in synaptic and neuronal dysfunction due to disruption of the proteostasis machinery [21, 25, 39, 71, 79].

Multiple pathways regulate protein synthesis at the level of translation initiation factor eIF2α. The integrated stress response (ISR) is a signaling network controlling translation initiation under stress, that enables cells to adapt to diverse homeostatic perturbations [19]. Four different kinases phosphorylate eIF2α at serine 51, blocking protein synthesis including PERK (PKR-like ER kinase), GCN2 (general control nonderepressible 2), PKR (double-stranded RNA dependent kinase) and HRI (heme-regulated inhibitor) [19]. eIF2α phosphorylation represses the action of GTP exchange factor eIF2B by turning eIF2 into a non-competitive inhibitor, which allows the translation of mRNAs containing upstream short reading frames (uORFs), highlighting the mRNA encoding activating of transcription factor 4 (ATF4) [85]. ATF4 controls the expression of multiple genes involved in proteostasis control, redox balance, autophagy, amino acid synthesis and mitochondrial fitness [3, 88]. Under prolonged eIF2α phosphorylation, ATF4 triggers apoptosis by engaging the proapoptotic factor CHOP, inducing the expression of members of the BCL-2 protein family, enhancing ROS production and reestablishing protein synthesis in stressed cells [88].

Studies in models of prion disorders, Parkinsońs disease and frontotemporal dementia with parkinsonism linked to chromosome 17 (FTDP-17) caused by *MAPT* mutations demonstrated that chronic phosphorylation of eIF2α negatively affects the synthesis of synaptic proteins, compromising brain function and neuronal plasticity [37, 51, 55, 56]. Importantly, cellular stressors that trigger the ISR enhance RAN translation in neuronal cultures and fly models (see examples in [15, 31, 33, 74, 90]), suggesting a feed-forward loop between DPR production and the stress generated in the cell. Among the different inducers of the ISR, ER stress is reported to play a relevant role in RAN translation at C9ORF72 repeat expansions [15, 33, 77, 90]. Other reports suggest that expanded antisense C4G2 mRNA also engages the ISR [27]. Studies on FTD tissue from patients carrying C9ORF72 mutations indicated high levels of ER stress, associated with the phosphorylation of PERK and eIF2α [27, 60]. Conversely, expanded DPRs have been shown to alter ribosome function [45, 50, 87], reduce translation [42, 53, 94], and alter stress granule dynamics [66, 78, 94], a phenotype that may involve physical interactions between DRPs and ribosomes [38, 49, 50, 53]. Overall, multiple studies suggest that the translation machinery plays a critical role in C9ALS/FTD disease pathogenesis.

Strategies to reduce eIF2α phosphorylation may have therapeutic applications in the context of neurodegenerative diseases [7, 40]. The oral administration of the PERK inhibitor GSK2606414 provides neuroprotection in multiple models of neurodegeneration including Tau-mediated FTD [55, 64], Parkinsońs disease [51], Huntingtońs disease [26], in addition to brain ischemia [23]. However, GSK2606414 has serious side effects, provoking pancreatic toxicity [34, 55]. On the other hand, ISRIB was identified as a highly selective drug to inhibit the consequences of eIF2α phosphorylation and thus to block ATF4 expression through the stabilization of eIF2B [69, 73, 81, 96]. ISRIB administration reduces the toxicity of poly(PR_20_) in cell culture models [44], in addition to providing protection in models of neurodegeneration [1], including cellular and fly models of C9ORF72 pathogenesis [15, 90, 93]. However, ISRIB shows poor solubility, hence limiting its translational potential. To overcome these issues, a drug screen of a NINDS library resulted in the discovery of the antidepressant trazodone and dibenzoylmethane (DBM) as alternative small molecules that block the ISR [35]. Administration of DBM or trazodone to Tau transgenic mice or prion-infected animals restored protein synthesis and provided remarkable neuroprotection [35].

Here we tested the possible beneficial effects of administering DBM to a viral-based model of C9ALS/FTD [16]. We found that the oral administration of DBM was sufficient to prevent cognitive impairment induced by the expression of C9ORF72 repeat expansion in mice. However, treatment with DBM did not reduce the content of poly(GA) and poly(GR) inclusions in the hippocampus and brain cortex. Finally, proteomic analysis of hippocampal tissue indicated that the main pathways altered by DBM in the C9ORF72 repeat expansion model were related to cytoskeleton organization, ALS and metabolic processes. Overall, our results suggest a protective effect of DBM administration on a model of C9ORF72 pathogenesis, reinforcing the concept that targeting protein synthesis may alleviate cognitive symptoms.

## Materials and methods

### Cell culture experiments

Neuro-2a (N2a) cells (CCL-131™ ATCC) were cultured in 24-well plates (1.8 x10^7^ cells/well) in Dulbecco’s Modified Eagle Medium (11965092, Gibco^TM^) supplemented with 10% heat-inactivated fetal bovine serum (16140071, Gibco^TM^) and 1% penicillin-streptomycin (15070063, Gibco^TM^). Cells were maintained in a humidified incubator at 37°C with 5% CO_2_. Cells were pretreated for 2 h with 50 μM of DBM followed by treatment with tunicamycin (654380, Calbiochem EMB Biosciences) at a final concentration of 500 ng/mL. ISR activation was measured using qPCR and western blot analysis using standard methods [84].

Human induced pluripotent stem cells (iPSC) from a healthy male (WTC11) were differentiated into neurons by inducible neurogenin-2 (NGN2) expression and co-cultured with rat astrocytes as described previously [61]. For the ER stress and puromycin experiments, 6-week-old human neurons were treated with vehicle (DMSO), thapsigargin (1 µM), or DBM for 2 hours followed by thapsigargin (with DBM) treatment. After treatments, neurons were washed once with fresh cell culture media then incubated with 10 µM puromycin for 15 minutes at 37 °C. Neurons were washed once then fixed for 15 minutes in 4% paraformaldehyde. Puromycin was visualized with a mouse primary antibody against puromycin (Kerafast, EQ0001) and an anti-mouse Alexa Fluor 488 secondary antibody (Thermo Fisher, A11029). Neurons were identified with an antibody against MAP2 (Cell Signaling, 4542S) and an anti-mouse Alexa Fluor 488 secondary antibody (Thermo Fisher, A21245). Images were acquired on a Nikon AX laser scanning confocal microscope. Western blots were performed as previously reported [84]. Image acquisition and analyses were performed blind to the experimental conditions.

### C9ORF72 repeat expansion model

Experiments were carried out using C57BL/ 6J wild-type mice. Animals were kept in cycles of 12:12 hours light / dark, at an ambient temperature of 22 ± 2 °C, with free access to food and water. Animal care and experiments were performed according to procedures approved by the Bioethics Committee of the Faculty of Medicine of the University of Chile (approved protocol CBA # 18214-MED-UCH).

Adeno-associated virus (AAV) packaged into serotype 9 type capsid, containing 2 or 66 repeats of G_4_C_2_ (AAV-2R and AAV-66R, respectively), were intracerebroventricular (ICV) injected in newborn C57BL/6J wild-type mice to achieve efficient transduction in neurons of the central nervous system [84]. Viral vectors for this study were generated as reported before [16]. Newborn mice were cryo-anesthetized on ice for a few minutes until no movements were observed. A 32-gauge needle (BD Biosciences) was inserted at a 30-degree angle, between lambda and the eye of the pup, and sunk to a depth of approximately two millimeters. Four microliters with 0.02% Fast Green (1×10^5^genomes/μl) of each virus were manually injected in the corresponding pups in the cerebral ventricle. Finally, pups were placed on a heat pad until they recovered movements and placed back in their cages. Animals were weaned at 21 days of age and then subjected to cognitive and motor studies at 3 and 6 months of age. Finally, animals were euthanized at 8 months for biochemical and histopathological analyses.

### DBM administration

Powdered chow was prepared by grinding standard food pellets and sieving the material to remove large particles. The resulting powder was mixed with DBM (Sigma) by vigorous manual agitation in a sealed plastic bag for 5 min, including shaking, inverting, and rotating, to ensure homogeneous distribution of the compound. The supplemented chow was then placed in a powder feeder jar within the cage (Braintree Scientific, Inc.).

To assess target engagement of DBM, wild-type mice were fed *ad libitum* with powdered chow containing 0.5% (w/w) DBM or vehicle (powdered chow) for 7 days. Tunicamycin was injected at a concentration of 50 μg/g using intraperitoneal injections and then tissue collected after 24 h as previously reported [11]. For long-term treatment, mice injected with AAV-2R or AAV-66R were fed ad libitum with powdered chow containing 0.5% DBM or vehicle starting at 1 month of age and continuing until the end of the experiment.

### Behavioral tests

To determine the progression of disease features of C9FTD/ALS mice, a battery of motor and behavioral tests was performed in the same experimental group. This battery includes open field assay, novel object recognition test (NOR), novel object location test (NOL), rotarod test and hanging wire test. All behavioral equipment was cleaned with 35% ethanol between each mouse. 14–16 mice were used per group to obtain adequate statistical power to detect changes in behavior. Female and male mice were included in the analysis.

*Open Field Test-* Open Field Test was used to measure cognitive deficits of the mouse. The apparatus consists of a 40 x 40 cm square box of polymethylmethacrylate (PMMA) with a video recorder on the top. Mice were placed in the center of the box and could freely explore the arena for fifteen minutes. Mice were recorded and movements were tracked and analyzed by the AnyMaze program (Stoelting, Co.) This program divides the box into 16 equal quadrants, of which 4 form the center, 4 the corners and a total of 8 form the periphery. This test was performed to evaluate anxiety behaviors. The distance traveled in the center zone versus the total distance traveled were plotted.

*Novel Object Recognition (NOR)-* The NOR task was used to assess recognition memory. NOR was performed in two consecutive days: on the first day (acquisition phase), mice were allowed to freely explore two identical objects for 5 min or until a cumulative exploration time of 20 s was reached. Exploration time for each object was recorded. On the second day (testing phase), 24 h later, one of the identical objects was replaced for a novel object, and interaction time with both objects was recorded. The percentage of time spent on the novel object was calculated using the following formula: (t new object/(t new object + t old object))*100%. Object pairs were randomly assigned to each animal during day 1, and the identity and spatial location of the novel object were counterbalanced across animals to minimize object and location preference biases. Animals that did not interact with any object (0%) or those that interacted with only one object (100%), were excluded from the analysis as reported (Leger et al., 2013; Vogel-Ciernia & Wood, 2014).

*Novel location recognition (NOL) –* NOL test was used to evaluate hippocampus-dependent long-term spatial memory in mice. A training test protocol similar to NOR was used. Unlike NOR, in the NOL test the location of one of the identical objects is changed on the second day of the test. The percentage of time spent exploring the novel object was calculated using the following formula: (t new location object/(t new location object + t old object))*100%.

*Hanging wire test-* Muscle strength and coordination were evaluated using the wire hang test. Mice were placed at the center of a horizontally stretched metal wire positioned 35 cm above the surface. The number of falls was recorded during a 2 min trial.

*Rotarod test –*Rotarod protocol was applied as was previously described [16]. Briefly, the rotarod test was performed over four consecutive days. On day 1, mice were briefly trained to familiarize them with the apparatus. Training was conducted at a constant basal speed of 4 rpm until the animal was able to maintain its grip on the rotating cylinder. Then, the test was performed by accelerating the rotation speed from 4 to 40 rpm over a 5 min period, and the latency to fall was recorded. Three trials per mouse were performed each day.

### Tissue preparation

Mice were euthanized by CO_2_ narcosis and perfused transcranial with 0.9% NaCl solution. Brains were removed and split in two hemispheres, where the right side was used for biochemical analysis and the left side for histopathology. Brain tissue was fixed in 4% PFA for histological analyzes to determine DPR inclusions, neuronal loss, and gliosis. Brain tissue was dehydrated through a battery of alcohols (two hours in alcohol at 80, 90, 95, 95, 100 and 100% each one), xylene (six hours) and the end, brains were put in paraffin (six hours) before being included in paraffin. Then 5 µm thick sections were serially cut on a microtome (microm HM 325, Thermo Fisher scientific) for analysis.

### Immunohistochemistry

The slides containing the paraffin-embedded sections were deparaffinized and rehydrated in xylene (2 times, 5 minutes each) and a battery of alcohols (100, 95, 90, 80, 70%) for 1 minute at each concentration and then washed in water (1 min). The endogenous peroxidase was then blocked with a 0.3% H_2_O_2_ solution (Merck, Germany) in PBS solution. After that, slides were washed in PBS (3 times, 5 minutes each) and antigen retrieval was performed in a steam chamber (30 min) with citrate buffer (pH 6.0). After washing twice with PBS, sections were blocked using a blocking solution (5% BSA, 0.3% Triton in PBS) (45 minutes) or the M.O.M. kit (mouse-on-mouse) (BMK-2202, Vectastain) for 1 h depending on primary antibody host. Finally, sections were incubated at room temperature in a wet chamber overnight, with the corresponding primary antibody. The primary antibodies used were: anti-NeuN (1: 400) (# MAB377, Millipore); anti-Poly (GA) (1: 1500) (# TIP-C9-P01, CosmoBio); anti-Poly (GR) (1: 32000) (# TIP-C9-P01, CosmoBio); anti-GFAP (1: 200) (# ab7260, Abcam), anti-Iba1 (1: 500) (# 019-19741, Wako).

### Gliosis

To quantify activation of microglia and astrocytes, immunostainings were performed with antibodies for Iba1 and GFAP, respectively. Images were taken in the hippocampus with an optical microscope (Leica DM500) at a magnification of 10X. Two sections were taken per mouse to have a technical duplicate. The quantification was performed in the hippocampal region using ImageJ software (NIH, Bethesda, United States) by determining the percentage of positive signal per each area.

### DPR inclusions: poly(GA) and poly(GR)

In order to quantify DPR expression, immunohistochemistry of poly(GA) and poly(GR) was performed. To this purpose, images of cortex (motor and somatosensory), hippocampus (CA1) and cerebellum were taken with an optical microscope (Leica DM500) at a magnification of 20X. Two sections per mouse were used as a technical duplicate. Quantification was performed manually, counting the positive cells for the signal of both poly dipeptides.

### Neuronal quantification

Anti-NeuN immunohistochemistry was performed to quantify neurons. The entire motor and somatosensory cortex was digitally acquired in an optical microscope (Leica DM500) at 10X magnification. Two sections were taken per mouse to obtain a technical duplicate. The quantifications were carried out using the ImageJ software (NIH, Bethesda, United States).

### RNA extraction and RT-PCR

RNA was extracted from the cortex, cerebellum, and hippocampus. Tissues were homogenized in TEN buffer solution (10mM Tris-HCl, 1mM EDTA, 100mM NaCl, pH 8.0) with protease and phosphatase inhibitors (Sigma Aldrich). Then, 50 µl of the homogenate was mixed with the TRIzol reagent (Life technologies) to extract RNA according to the manufacturer’s instructions. The remaining volume was stored at-80 °C as a backup. Following this, 1 µg of RNA was used to generate cDNA by a reverse transcriptase polymerase chain reaction (RT-PCR) using a high-capacity reverse transcription kit (# 4368813, Applied Biosystems) according to manufacturer’s instructions.

Real time PCR reactions were performed using the EvaGreen reagent (Biodyne, United States) following the supplier’s instructions in a Stratagene Mx3000P system (Agilent Technologies). The thermal profile used for qPCR was: 1 denaturation cycle of 95 °C for 10 min; 40 cycles of amplification of 95 °C for 10 s, 58 °C for 15 s, 72 °C for 20 s; 1 final amplification cycle of 95 °C for 15 s, 25 °C for 1 s, 70 °C for 15 s and 95 °C for 1 s. Relative amounts of mRNA were calculated from comparative threshold values of 40 cycles using Actin mRNA as internal expression control. All the quantifications were plotted as mRNA relative levels. The values were calculated using the method “ΔCt”, determining the power in base 2 of the delta between the housekeeping gene and gene of interest by the following formula: ΔCt= 2(Ct (housekeeping gene) - Ct (gene of interest)). All primers are described in Additional file 1: Table S1.

### Proteomic study and bioinformatic analysis

*Proteomic Sample Preparation.* Hippocampi dissected from mouse brains were frozen on dry ice and stored at-80 °C. Tissues were suspended in 150 µL of TEN buffer (10 mM Tris-HCl, pH 8; 1 mM EDTA, pH 8; 100 mM NaCl with protease and phosphatase inhibitors) and homogenized by grinding with a mortar and pestle on ice. 100 µg protein aliquots were set aside and digested using micro S-Trap columns (Protifi).

*Digestion.* Homogenized tissue samples were added to a lysis buffer containing a final concentration of 5% SDS and triethylammonium bicarbonate (TEAB), pH ∼7.55. Samples were reduced in 20 mM dithiothreitol (DTT) for 10 minutes at 50 ⁰C with agitation, then cooled at room temperature for 10 minutes, and finally alkylated with 40 mM iodoacetamide (IAA) for 30 minutes at room temperature in the dark. Samples were acidified with a final concentration of 1.2% phosphoric acid, resulting in a visible protein colloid. S-Trap binding buffer (90% methanol in 100 mM TEAB) was added at a volume of 6 times the acidified lysate volume. Samples were vortexed until the protein colloid was completely dissolved in the 90% methanol. The entire volume of the samples was spun through the micro S-Trap columns in a flow-through microcentrifuge tube. Samples were spun through in 200 µL aliquots for 10 seconds at 4,000 x g. Subsequently, the S-Trap columns were washed with 200 µL of the S-Trap binding buffer twice for 10 seconds each at 4,000 x g. S-Trap columns were placed in a clean elution tube and incubated at 47⁰ C for 1 hour with 125 µL of trypsin digestion buffer (50 mM TEAB, pH ∼8) at a 1:25 ratio (protease:protein, wt:wt). The same trypsin digestion buffer was added a second time for an incubation at 37⁰ C overnight. Peptides were eluted from the S-Trap column the next morning in the same elution tube as follows: 80 µL of 50 mM TEAB was spun through for 1 minute at 1,000 x g. 80 µL of 0.5% formic acid was spun through next for 1 minute at 1,000 x g. Finally, 80 µL of 50% acetonitrile in 0.5% formic acid was spun through the S-Trap column for 1 minute at 4,000 x g. These pooled elution solutions were dried in a vacuum concentrator and then re-suspended in 0.2% formic acid.

*Desalting.* The re-suspended peptide samples were desalted with stage tips made in-house containing a C18 disk, concentrated and re-suspended in aqueous 0.2% formic acid containing “Hyper Reaction Monitoring” indexed retention time peptide standards (iRT, Biognosys).

*Mass Spectrometric Analysis.* LC-MS/MS analyses were performed on an Eksigent Ultra Plus nano-LC 2D HPLC system combined with a cHiPLC system directly connected to an orthogonal quadrupole time-of-flight (Q-TOF) SCIEX TripleTOF 6600 mass spectrometer (SCIEX). The solvent system consisted of 2% ACN, 0.1% FA in H_2_O (solvent A) and 98% ACN, 0.1% FA in H_2_O (solvent B). Proteolytic peptides were loaded onto a C_18_ pre-column chip (200 μm × 6 mm ChromXP C18-CL chip, 3 μm, 300 Å; SCIEX) and washed at 1 μL/minute for 10 minutes with the loading solvent (H_2_O/0.1% FA) for desalting. Peptides were transferred to the 75 μm × 15 cm ChromXP C_18_-CL chip, 3 μm, 300 Å (SCIEX) and eluted at 300 nL/min with the following gradient of solvent B: 5% for 5 min, linear from 5% to 8% in 15 min, linear from 8% to 35% in 97 min, up to 80% in 20 min, 80% for 10 min, and back to 5% in 3 min, The column was re-equilibrated for 30 min with 5% of solvent B, and the total gradient length was 180 min. All samples were analyzed by data-independent acquisition (DIA), specifically using variable window DIA acquisitions [18, 30, 68]. In these DIA acquisitions, 64 windows of variable width (5.9 to 90.9 m/z) are passed in incremental steps over the full mass range (m/z 400–1,250), as determined using the SWATH Variable Window Assay Calculator from SCIEX (Additional file 1: Table S2). The total cycle time of 3.2 seconds includes a MS1 precursor ion scan (250 msec accumulation time), followed by 64 variable window DIA MS/MS segments (45 msec accumulation time for each). MS2 spectra were collected in “high-sensitivity” mode. The collision energy (CE) for each segment was based on the z=2+ precursor ion centered within the window with a CE spread of 10 or 15 eV.

*DIA Data Processing and Statistical Analysis.* All DIA data were processed in Spectronaut version 14.2.200619.47784 (Biognosys) using an in-house custom-made mouse brain spectral library (32,334 modified peptides and 3,246 protein groups). Data extraction parameters were selected as dynamic, and non-linear iRT calibration with precision iRT was selected. Identification was performed using a 1% precursor and protein q-value, and iRT profiling was selected. Quantification was based on the MS/MS peak area of the 3-6 best fragment ions per precursor ion, peptide abundances were obtained by summing precursor abundances and protein abundances. Interference correction was selected, and local normalization was applied. Differential protein abundance analysis was performed using paired t-test, and p-values were corrected for multiple testing, specifically applying group-wise testing corrections using the Storey method [9, 76]. For the differential analysis, protein groups with at least two unique peptides and q-value ≤ 0.05 were significantly altered (Additional file 1: Table S2).

Hippocampal tissue samples (5 female mice per condition) were homogenized in TEN buffer containing protease and phosphatase inhibitor cocktail (Roche). Protein samples were digested and TMT labeled as we reported [82]. Liquid chromatography-tandem mass spectrometry analysis was performed using an Exploris 480 mass spectrometer equipped with an Ultimate 3000 RSLCnano system (ThermoFisher). The global correlation analysis was performed using log2 fold-changes from the differential abundance analysis. Enrichment analysis was calculated using the R package STRINGdb 2.4.2 with the STRING database version 1184. Protein interaction networks were generated using the stringApp in Cytoscape 3.8.285 and nodes were colored according to statistically significant enriched biological processes and cellular components.

### Transcriptomic data acquisition and pre-processing

Transcriptomic analyses were conducted in the R statistical environment (v4.4.1). Raw RNA-Seq count data from the GSE137810 dataset were retrieved via the GEOquery package. The transcriptomic datasets analyzed during the current study are available in the Gene Expression Omnibus (GEO) repository under the accession number GSE137810 [https://www.ncbi.nlm.nih.gov/geo/query/acc.cgi?acc=GSE137810]. To mitigate sequencing depth heterogeneity, raw counts were normalized to Counts Per Million (CPM). Batch effects were rigorously corrected using the ComBat algorithm (sva package), modeling the diagnosis (ALS/FTD vs. CTRL) as the primary biological covariate to preserve disease-associated variance while removing technical noise. Anatomical annotations were standardized to consolidate spinal cord segments into a unified analytical category. To assess relative expression changes of candidate genes (INA, TUBB4B, VIM, CADPS, GDI1, NCDN) across heterogeneous tissues, expression values were standardized to Z-scores. Pairwise comparisons between ALS/FTD and Control groups were performed using unpaired t-tests, with statistical significance thresholds set at *p* < 0.05. Transcriptional modulation of stress mechanisms was evaluated using two complementary approaches. First, Gene Set Enrichment Analysis (GSEA) was performed on pre-ranked gene lists (sorted by Log2 Fold Change) using the clusterProfiler package to calculate Normalized Enrichment Scores (NES) for the’Integrated stress response signaling’ (GO:0140467) and’BIOCARTA_EIF2_PATHWAY’ (Biocarta M6924). Second, to resolve inter-individual variability, Gene Set Variation Analysis (GSVA) was employed using the GSVA package with Gaussian kernels. This yielded single-sample enrichment scores for the ISR pathway, enabling downstream correlation analyses. Functional clustering was further refined by calculating semantic similarity using GOSemSim to dissect the ISR gene set into functional modules based on Gene Ontology topology.

## Statistical analyses

Statistical analyses were performed using Prism V7 software (GraphPad Software Inc., USA). Data were analyzed using one-way ANOVA and Tukey’s post hoc test for multiple variables. All data in bar graphs show mean ± standard deviation (SD). For bioinformatic analysis, correlations between pathway activity scores (GSVA) and gene expression (LogCPM) were assessed using Pearson’s correlation coefficients. All visualizations were generated using ggplot2, with statistical annotations derived from the tests. Statistical differences were considered significant for values of *p* < 0.05. The p-values are displayed as follows: \**p* < 0.05, \*\**p* < 0.01 and \*\*\**p* < 0.001.

## Results

### DBM reduces the activity of the ISR

To validate the activity of DBM on the ISR, we treated Neuro2A cells with 50 μM DBM for 2 h, followed by the exposure to 500 nM of the ER stress agent thapsigargin, a SERCA inhibitor. After 8 h, ATF4 protein levels were determined by Western blot (Fig. 1b). Treatment with DBM reduced the upregulation of ATF4 (Fig 1b). An unexpected modulation of basal ATF4 levels by DBM was observed in this experiment. The mRNA levels of three ATF4 target genes, including *Atf3*, *Chop* and *Gadd34* were determined using qPCR [51]. DBM administration significantly reduced the levels of ATF4-target genes in Neuro2a cells (Fig. 1c). Next, new protein synthesis was measured in human iPSC-derived neurons. Cells were treated with thapsigargin for 2 hours to induce an ER stress response and protein synthesis was monitored by measuring puromycin incorporation into newly synthesized proteins. Thapsigargin treatment significantly reduced protein synthesis in the soma of human neurons, an effect that was abolished by DBM treatment (Fig. 1d).

**Figure 1.**
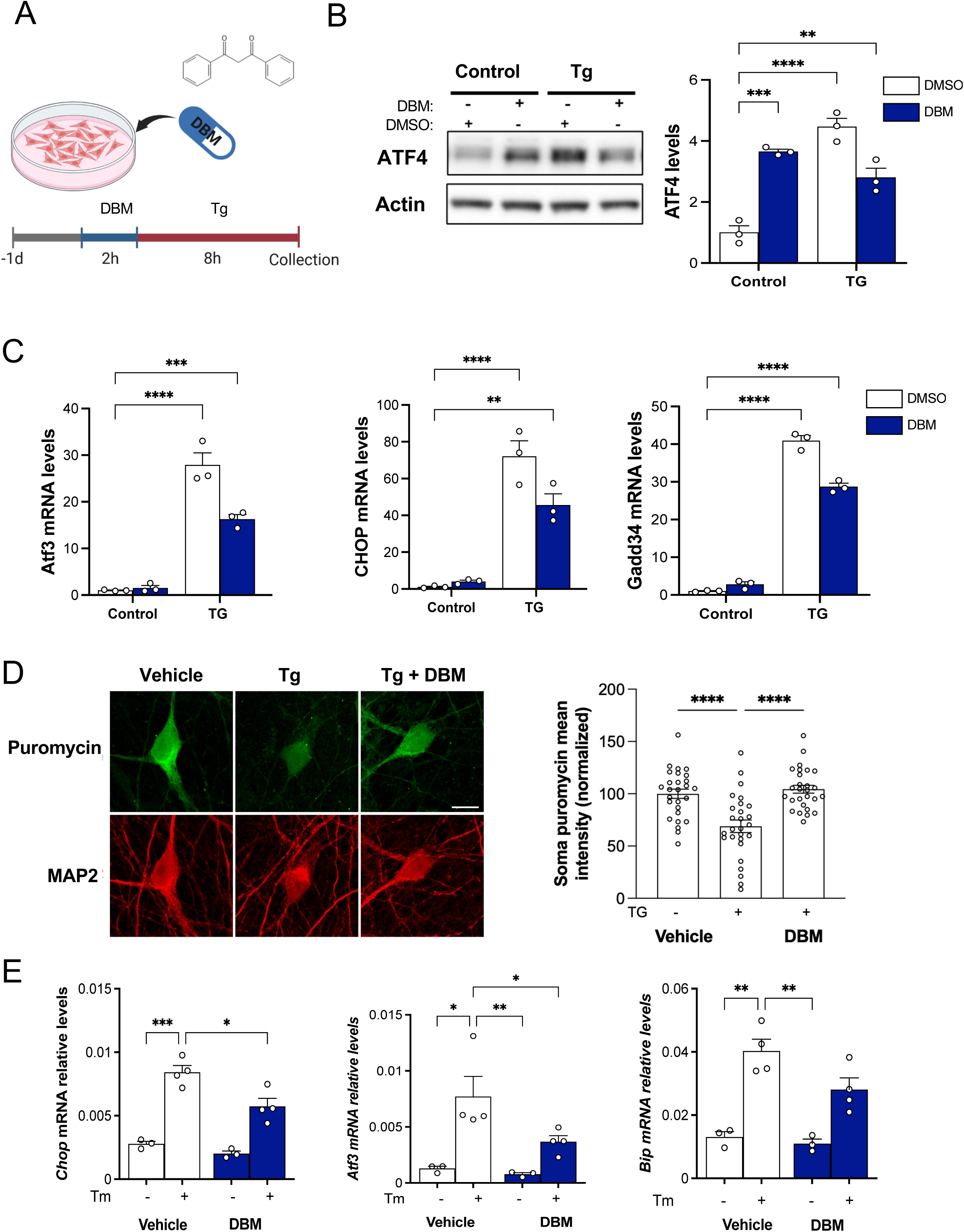
DBM administration reduces the ER stress response. **(A)** Experimental design: Neuro2A cells were pre-incubated for 2 h with either 50 µM DBM or vehicle control (DMSO) and subsequently treated with 500 nM thapsigargin (Tg) or vehicle for additional 8 h. Cells were then harvested, and total protein and RNA were extracted. **(B)** ATF4 protein abundance was evaluated by western blot analysis and normalized to β-actin abundance. **(C)** Relative mRNA expression levels of downstream ATF4-responsive genes *Atf3*, *Chop*, and *Gadd34* were quantified by qRT-PCR and normalized to β-actin transcript. In B and C, quantitative data represent means ± SEM from three independent experiments and were subjected to analysis of variance (one-way ANOVA; *: *p* < 0.05, ****: *p* < 0.0001). **(D)** Representative images of immunostaining against puromycin-labeled newly synthesized proteins (green) and MAP2 (red) from human iPSC-derived neurons treated with vehicle, Tg, or Tg with DBM. Scale bar: 10 µm. Right panel: quantification of the normalized mean puromycin intensity in the neuronal soma of 3 independent experiments (n = 27 / group, ***: *p* < 0.001, 1-way ANOVA, Bonferroni post-hoc analyses). Each dot represents one cell. **(E)** One-month-old mice were fed ad libitum with powdered chow containing 0.5% dibenzoylmethane (DBM) or vehicle for 7 days. Animals were then injected with tunicamycin (Tm), and brain tissue was collected 24 h later. qPCR analysis was performed in total cDNA to measure *Chop*, *Atf3*, and *Bip* mRNA levels in frontal cortex of animals treated with 0.5% DBM or vehicle, followed by injection of Tm. All data were normalized with actin mRNA levels. Data are presented as mean ± S.D. *: *p* < 0.05. **: *p* < 0.01, ***: *p* < 0.001. Two-way ANOVA and Dunnett’s multiple comparisons test.

To assess the activity of DBM in the CNS through oral administration, we used a pharmacological model of ER stress based on the intraperitoneal injection of tunicamycin (N-glycosylation inhibitor), which induces the activation of the UPR in the CNS [11]. Mice were pre-treated for 1 week with DBM by mixing the compound with powdered food (0.5% *ad libitum*) and then animals were injected with 5 µg of tunicamycin per gram of mice. After 24 h, brain cortex was dissected and mRNA expression levels of the ATF4-target genes *Chop, Atf3,* and *Bip* were measured by qPCR (Fig. 1e). Remarkably, the expression levels of these three ER stress-responsive genes were decreased in the frontal cortex of animals pretreated with DBM compared with control animals following tunicamycin injection (Fig. 1e). We also measured the levels of *Xbp1* mRNA splicing, a parallel signaling pathway triggered by ER stress. No significant changes were observed in *Xbp1s* transcript level in animals pretreated with DBM following tunicamycin-induced ER stress in frontal cortex, although significant reduced levels were found in hippocampus (Fig. S1).

### DBM administration prevents memory impairment on a mouse model of C9ORF72-repeat expansion

Several mouse models have been developed to study C9ORF72 pathogenesis, showing phenotypic heterogeneity ranging from a lack of disease manifestation, low penetrance, high phenotypic variability over a large range of time, sex dependency, and even differences generated by the animal housing conditions due to differential intestinal microbiota [8, 52, 57]. To study C9ORF72 repeat expansion pathogenesis, we used an AAV-based model previously employed in our laboratory that consisted of the overexpression of 66 GGGGCC repeats (AAV-66R) in the CNS by injecting viral particles into the ventricle of newborn pups. This animal model recapitulates several ALS/FTD features [16]. We generated four experimental groups defined as AAV-2R/vehicle (n = 14), AAV-66R/vehicle (n = 17), AAV-2R/DBM (n = 17), and AAV-66R/DBM (n = 15). Mice were subjected to different motor and cognitive tests including open field test, novel object recognition (NOR), novel object location (NOL), hanging wire test and rotarod at 3 and 6 months of age.

NOR and NOL evaluate the capacity of animals to discriminate between familiar and novel objects or the displacement of objects in different quadrants of the cage, respectively (Fig 2a and 2c). The NOR test has been widely used in assessing non-spatial object memory in rodents, which is dependent on the perirhinal cortex and hippocampus; whereas NOL provides information about spatial memory acquisition and is associated with hippocampus-dependent memory formation. At 3 months old, the viral model did not develop clear cognitive impairment using these two tests. However, at 6 months of age, AAV-66R mice showed disrupted cognitive performance when compared to AAV-2R using NOR and NOL (Fig. 2). Remarkably, treatment with DBM resulted in almost complete protection in AAV-66R-injected mice as assessed with the NOR assay, showing behavioral performance similar to control animals (Fig. 2b). Virtually identical results were obtained when the NOL test was assessed (Fig. 2d), suggesting important cognitive improvement upon DBM administration. To evaluate anxiety-like behavior, the open field test was also performed. However, no significant differences were found between animals injected with AAV-66R or control vector at 3 or 6 months of age (Additional file 1: Fig. S2c).

**Figure 2.**
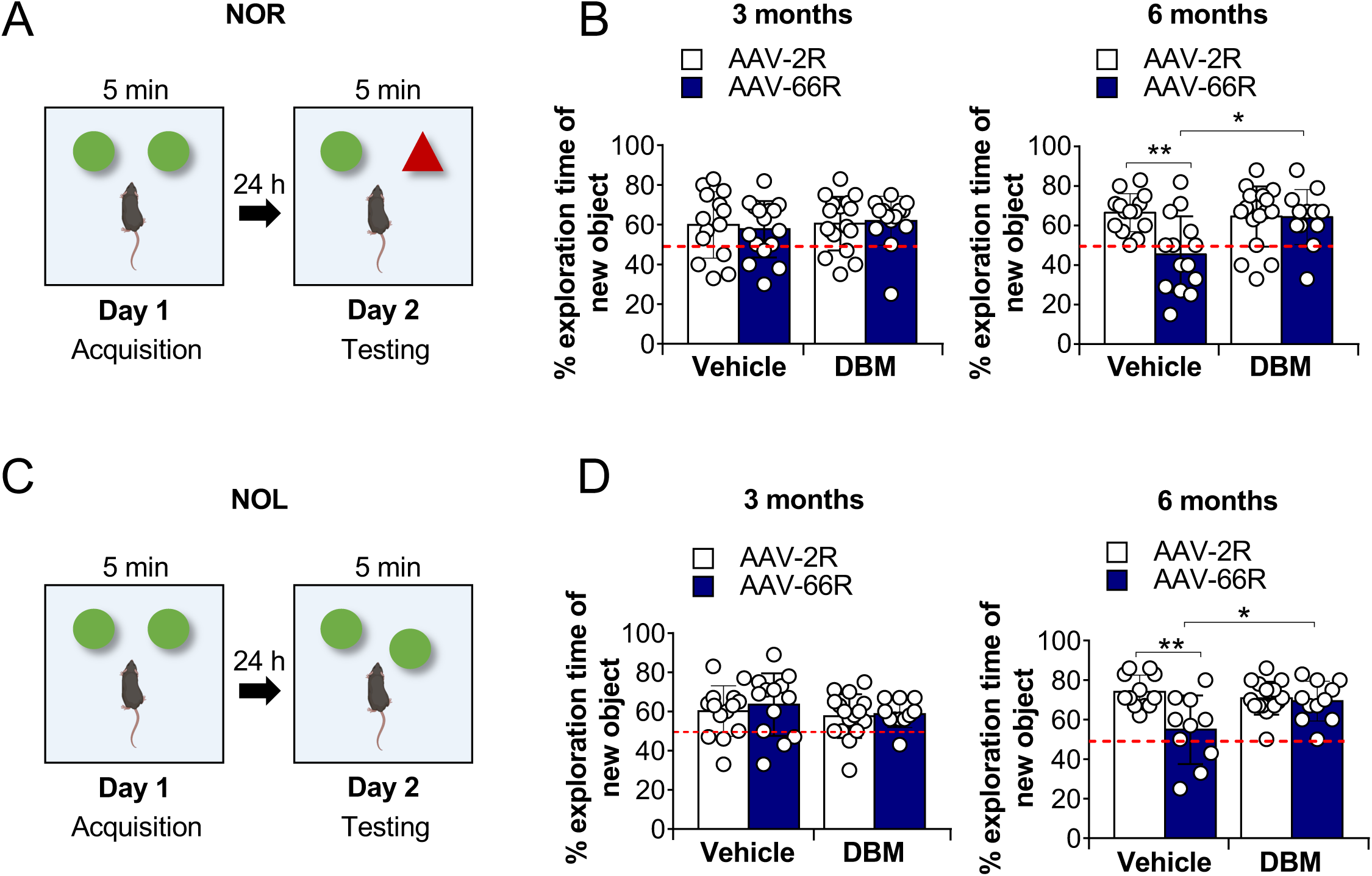
Oral administration of DBM prevents memory impairment on a C9ORF72-mediated FTD/ALS mouse model. Animals were injected with the AAV-2R or AAV-66R at P0-1 and fed with powdered chow containing 0.5% dibenzoylmethane (DBM) or vehicle *ad libitum* from the age of 1 month until the end of the experiment. **(A)** Schematic representation of the NOR test: on day 1, mice were exposed to two identical objects, and interaction time with each object was recorded for 5 min. Then, 24 h later, one object was replaced with a new one and, interaction time with each object was recorded for 5 min. **(B)** The NOR test was performed in animals at 3 and 6 months of age. The percentage of exploration time of the novel object on the second day of the test is shown. Dashed line indicates 50% of exploration time spent exploring the novel object. Values above 50% indicate a preference for the novel object. **(C)** Schematic representation of the NOL test: in day 1, mice were exposed to two identical objects and interaction time with each object was recorded for 5 minutes. Then, 24 hours later, one object was moved to a new location and interaction time with each object was recorded for 5 minutes. **(D)** NOL test was performed in animals at 3 and 6 months of age. The percentage of exploration time of the novel location on the second day of the test is shown. The dashed line indicates 50% of the time spent exploring the object in the novel position. Each dot represents one animal. Data are presented as mean ± S.D. (n = 14-17). *: *p* < 0.05. **: *p* < 0.01; one-way ANOVA followed by Tukey’s multiple comparisons test.

No clear motor impairment was detected in AAV-66R–injected mice, as assessed by the rotarod test (Additional file 1: Fig. S2a). Consistently, performance in the hanging wire test revealed no significant differences between groups when animals were evaluated at either 3 or 6 months of age (Additional file 1: Fig. S2b). Together, these findings indicate that, under the experimental conditions used in this study, the model predominantly recapitulates behavioral features associated with FTD rather than ALS.

We also monitored body weight as a measure of health status. Analysis by gender indicated that only male animals injected with AAV-66R showed a body weight decrease at six months of age (around 11% reduction), which was prevented by DBM treatment (Additional file 1: Fig. S3). Overall, our results suggest that C9ORF72 repeat expansion expression in mice triggers cognitive defects that are reversed by the oral administration of DBM.

### Histopathological alterations associated with G_4_C_2_ repeat expression are not affected by DBM administration

To assess the presence of DPR inclusions due to G_4_C_2_ repeat expansion expression, we performed immunohistochemistry to detect poly(GA) and poly(GR) DPR content in hippocampus and cortex. The total number of inclusion-positive cells in both tissues was quantified for each animal (Fig. 3A-D). A significant number of poly(GA) positive cells were found in both hippocampus (CA1 region) and cortex of AAV-66R mice. Nevertheless, no significant effects of DBM administration were observed when poly(GA)-positive cells were quantified (Fig. 3b). Similar results were obtained after quantifying the number of poly(GR)-positive cells (Fig. 3d). Additionally, DPR-positive cells were classified according to the cellular distribution of protein aggregates as perinuclear, cytoplasmic, or diffuse cytosolic patterns (Additional file 1: Fig. S4) and no significant differences were observed between DBM and vehicle treated animals (Additional file 1: Fig. S4). The number of DPR-positive cells in the cerebellum was very low; therefore, no further quantification was performed in this region.

**Figure 3.**
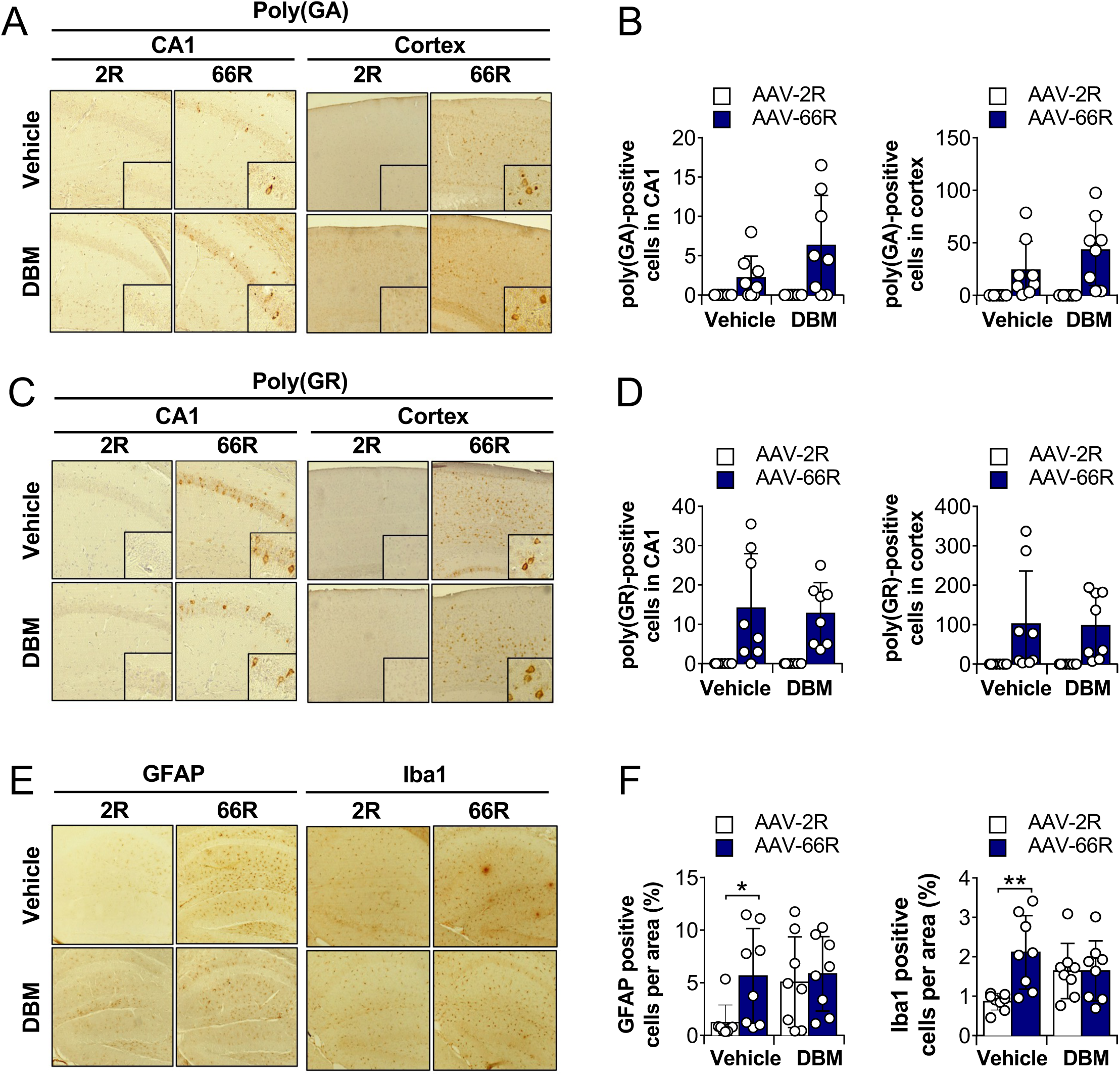
Histopathological characterization of animals injected with AAV-66R mice. Animals were injected with the AAV-2R (2R) or AAV-66R (66R)at P0-1 were fed with powdered chow containing 0.5% DBM or vehicle *ad libitum* from the age of 1 month until the end of the experiment. **(A)** Representative images of poly(GA)-labeled cells in the CA1 region (left) and cortex (right) of mice injected with AAV-2R or AAV-66R and treated with vehicle or DBM via oral administration. **(B)** Quantification of the number of poly(GA)-positive cells in the CA1 region (left) and cortex (right). **(C)** Representative images of poly(GR)-labeled cells in the CA1 region (left) and cortex (right) of mice injected with (2R) or (66R) and treated with vehicle or DBM via oral administration. **(D)** Quantification of the number of poly(GR)-positive cells in the CA1 region (left) and cortex (right). **(E)** Representative images of GFAP-labeled cells (left) and Iba1-labeled cells (right) in hippocampus of mice carrying (2R) or (66R) treated with vehicle or DBM. **(F)** Quantification of GFAP-positive (left) and Iba1-positive (right) cells in the hippocampus. Values are expressed as the percentage of positive signal per area. Each dot represents one animal (n = 8 per group, male and female mice). Data are presented as mean ± S.D. Unpaired-t test, *, *p* < 0.05. **; *p* < 0.01. Tissue was collected from animals at 6 months of age for histological analysis.

We also measured other histopathological markers associated with neurodegeneration, including gliosis and neuronal loss. Analysis of the microglial marker Iba1 using immunohistochemistry indicated enhanced microglial activation in the hippocampus and cortex of AAV-66R mice (Fig. 3e and 3f, right panels). Unexpectedly, DBM treatment enhanced the signal of Iba1 at basal levels, similar to the values obtained in the disease model. A similar pattern was observed when GFAP staining was performed as a measure of astrogliosis (Fig. 3e and 3f, left panels). Analysis of brain cortex indicated very low signals of gliosis in the disease model (data not shown). Finally, neuronal content in the brain cortex was measured after NeuN staining using immunohistochemistry. However, under the conditions tested, injections of AAV-66R did not result in neuronal loss in the brain cortex (Fig. S6). Taken together, our results suggest that DBM administration does not alter the accumulation of DPRs or gliosis in our C9ORF72 mouse model.

### Proteomics analysis of hippocampal tissue of C9ORF72 repeat expansion animals treated with DBM

To gain insights into the possible molecular mechanisms mediating the positive effects of DBM administration on the cognitive performance of our C9ALS/FTD model, we performed proteomic profiling by LC-MS/MS of hippocampus of all experimental groups. This region was analyzed because it is directly related to the behavioral assessments performed in this study. First, we defined the proteomic alterations triggered by the expression of DPRs at the whole proteome level according to fold change values, comparing AAV-66R-vehicle with AAV-2R-vehicle derived tissue, and then with AAV 66R-DBM treated animals. Interestingly, we found a global reversion of the proteome changes triggered by the disease model upon DBM administration with a negative Spearman correlation of 0.64, corresponding to 73% of whole identified proteins that expression levels were altered in opposite directions (Fig. 4a and Additional file 1: Table S2).

**Figure 4.**
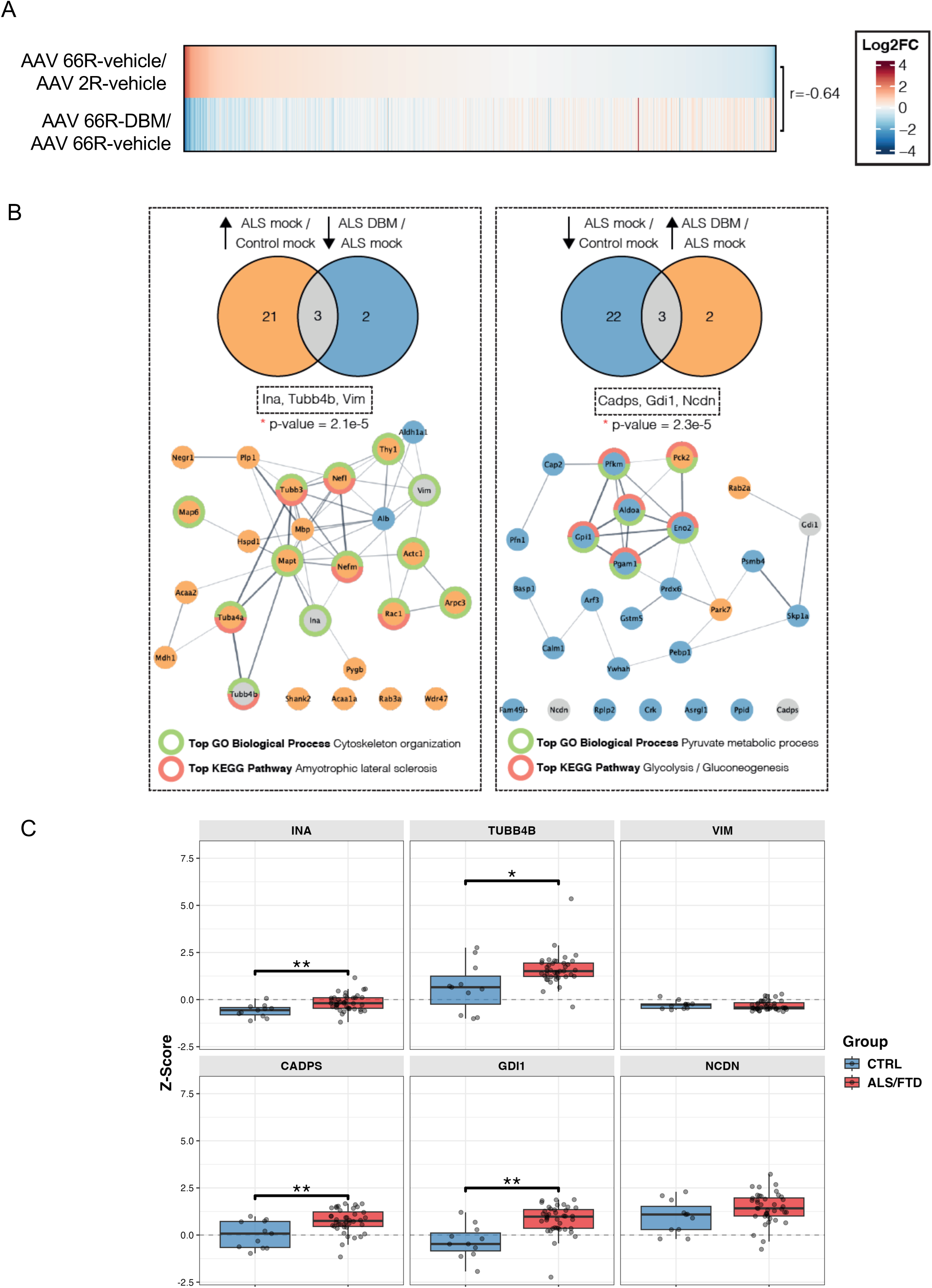
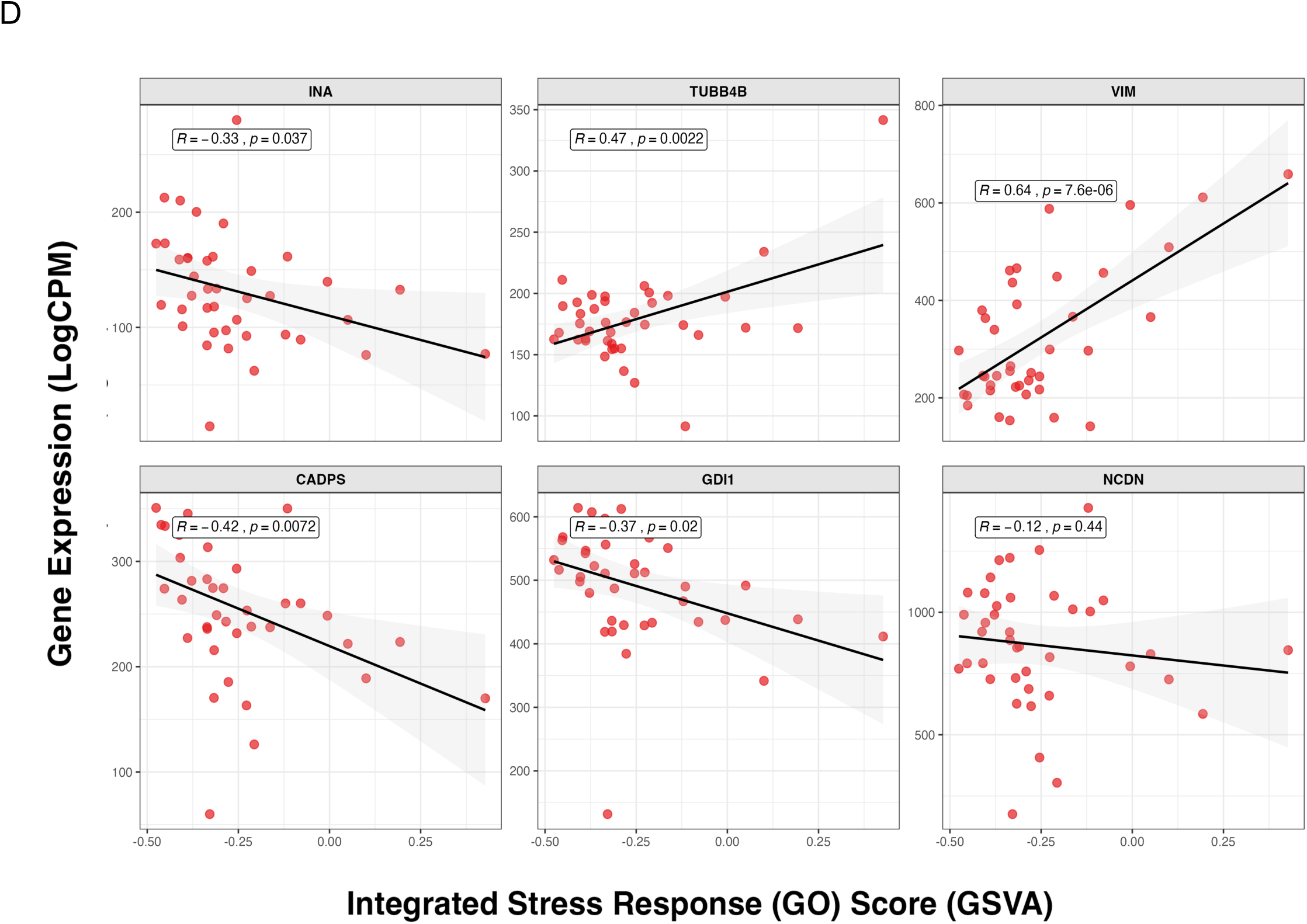
Differential protein expression in hippocampus of C9ORF72-mediated FTD/ALS mice identified by LC-MS/MS. Animals were injected with AAV-2R or AAV-66R at P0-1 and then fed with powdered chow containing 0.5% dibenzoylmethane (DBM) or vehicle ad libitum from the age of 1 months until the end of the experiment. Hippocampal tissue was collected from animals at 6 months of age for quantitative proteomics analysis. (n = 5 female per group). **(A)** Global correlation analysis of the proteomic changes observed in the differential abundance analysis (r= 0.64). At the right of the heatmap, r values represent Spearman’s correlation coefficient between the comparisons. **(B)** Upper panel: Venn diagram of overlaps between the proteins changing abundance in opposite directions in the disease model compared with controls and disease model treated with DBM. Bottom panel: Interaction network of proteins corrected by DBM treatment in C9ORF72 mice. **(C)** Differential expression of DBM-responsive genes identified in B now in human ALS/FTD post-mortem tissue (NYGC ALS Consortium dataset GSE137810). Z-score normalized expression of selected genes (*INA, TUBB4B, VIM, CADPS, GDI1, NCDN*) in the hippocampus of control (CTRL, n = 11) and ALS/FTD (n = 40) subjects. Center line: median; box limits: upper and lower quartiles; whiskers: 1.5x interquartile range. Significance was assessed via unpaired two-tailed t-test (*: *p* < 0.05, **: *p* < 0.01, ***: *P* < 0.001, ****: *p* < 0.0001). **(D)** Correlation between the presence of a gene expression signature of the ISR pathway and DBM-responsive genes. Scatter plots showing the GSVA enrichment score for the ISR pathway (x-axis) versus the log-transformed expression of selected markers (y-axis) in the hippocampus. Dots represent individual ALS/FTD samples. Black lines indicate linear regression with 95% confidence intervals (gray shading). Pearson’s *R* and *P*-values are shown.

Then, to identify specific proteins altered in the C9ORF72 model that are corrected by DBM (q<0.05), a Venn diagram analysis was used to display the overlap between these groups. We found six proteins that changed in opposite directions, where Alpha-internexin (INA), Tubulin Beta 4B (TUBB4B) and Vimentin (VIM) showed increased levels in the disease model but were downregulated by DBM administration (Fig. 4b). Calcium Dependent Secretion Activator (CADPS), GDP Dissociation Inhibitor 1 (GDI1) and Neurochondrin (NCDN) were downregulated in the C9ORF72 model but upregulated by DBM treatment. Additionally, protein-protein interaction network analysis showed most altered proteins belong to the same interaction cluster (Fig. 4b). Gene set enrichment analysis was performed to highlight main biological processes significantly affected in the disease model and by DBM (q < 0.05) (Fig. 4b). Notably, our analysis of upregulated genes reduced by DBM treatment indicated that the top enriched term in the gene ontology database was “Cytoskeleton organization”, in which 13 of the 24 proteins that were upregulated in the model (AAV 66R-vehicle/AAV 2R-vehicle) were associated to this cellular function. This analysis also showed that 6 of these 13 proteins are associated with ALS (top enriched terms in KEGG pathway), suggesting that the AAV-66R model may recapitulate, at the proteome level, some molecular alterations related to ALS pathogenesis.

Analysis of downregulated proteins (AAV 66R-vehicle versus AAV 2R-vehicle) that were upregulated by DBM administration indicated that the top enriched terms in the gene ontology database and KEGG pathway were “Pyruvate metabolic process” and “Glycolysis/Gluconeogenesis”. 5 out of 25 proteins were associated with both terms, suggesting a strong association with metabolic processes affected by the expression of DRP (Fig. 4b). Notably, Rab2a and Gdi1 (right panel Fig. 4a), proteins that are upregulated in AAV-66R mice by DBM, belong to the Rab subfamily of the small GTPases and dDENN domain, which plays an important role in C9ORF72 pathogenesis [70, 89], hence suggesting that DBM may modify the expression of relevant disease factors. Overall, our results indicate that our AAV-C9ORF72 repeat expansion model recapitulates distinct proteomic alterations observed in ALS/FTD, in addition to indicating putative biological pathways altered by G_4_C_2_ repeat expansion that are targeted by DBM treatment.

To assess the possible clinical relevance of our findings, we analyzed the GSE137810 transcriptomic dataset, which comprises a comprehensive cohort of ALS/FTD patients (spinal cord, n = 672; hippocampus, n = 40) and healthy controls (spinal cord, n = 124; hippocampus, n = 11). We evaluated whether the DBM-regulated genes identified in our mouse model were dysregulated in humans. Indeed, a significant subset of these genes showed altered expression in human ALS/FTD post-mortem hippocampal tissue (Fig. 4c), with comparable alterations observed in the spinal cord (Additional file 1: Fig. S7a). Next, we assessed markers of ISR activity using a 51-gene expression signature previously defined (GO:0140467). Correlation analysis revealed a significant association between ISR activity (GSVA) and the expression levels of 5 out of the 6 DBM-rescued genes (Fig. 4d). Although global analysis of the ISR gene set in the hippocampus and spinal cord did not reveal significant enrichment in human ALS/FTD tissues (Additional file 1: Fig. S7b), examination of individual gene expression levels revealed heterogeneity within the pathway. Nevertheless, distinct clusters displaying homogeneous activation patterns across tissues were identified (Additional file 1: Fig. S7c). Notably, key eIF2 regulators, such as *EIF2S, EIF2AK1, EIF2AK3* and *ATF4*, were consistently upregulated across ALS/FTD tissues. Furthermore, analysis of a cluster of genes representing an eIF2α signaling signature (gene set M6924) revealed a significant positive enrichment in the hippocampus, consistent with our previous observations (Additional file 1: Fig. S7d). Collectively, these gene expression studies demonstrate that a subset of genes rescued by DBM treatment in our C9ORF72 mouse model are dysregulated in human ALS/FTD patients, suggesting possible clinical relevance of our findings.

## Discussion

The development of new mouse models to study C9ORF72 pathogenesis is expected to accelerate the understanding of the molecular mechanisms and the future design of experimental treatments for ALS and FTD [5, 16, 44, 54, 86]. The assessment of small molecules to target the ISR in different neurodegenerative diseases has validated a possible strategy to treat protein misfolding disorders [34, 35, 73]. However, the therapeutic potential of the ISR in C9ORF72 awaited validation in mammalian *in vivo* models. A close relationship between the production of C9ORF72-derived DPRs and the activation of the ISR has been reported (see examples in [15, 33, 90, 93]). In cellular models of C9ORF72, the activation of the ISR promotes RAN translation, generating more DPRs and consequently further enhances eIF2α phosphorylation, sustaining a positive feed-forward loop between DPR production and eIF2α phosphorylation [15, 33, 90]. However, targeting the ISR with DBM did not reduce DRPs accumulation in contrast to previous studies using cell culture and fly models. It is possible that the partial effects of DBM on ISR inhibition were not sufficient to affect DRP aggregation. Importantly, oral administration of DBM improved the cognitive capacity of C9ORF72 mice as assessed with the NOR and NOL assays but did not reduce gliosis. However, DBM treatment had basal effects in wild-type mice that warrant future investigation. Together these results suggest that the beneficial effects of DBM are possibly linked to improved synaptic function, rather than affecting protein aggregate formation or their clearance.

Small molecules that target the ISR, such as ISRIB, reverse the effects of eIF2α phosphorylation by binding to eIF2B [73, 81, 96]. ISRIB has impressive neuroprotective effects in different mouse models of neurodegenerative diseases without causing pancreatic toxicity, including ALS [32, 34, 58, 91]. Additionally, ISRIB administration improves synaptic plasticity of wild-type mice [73] and reverses cognitive deficits after traumatic brain injury [17]. Cell culture experiments suggested that ISRIB treatment reduces RAN translation [15]. However, ISRIB has solubility problems, making it unsuitable for use in humans [34]. DBM, a natural product found in licorice root, mimics the effects of ISRIB. However, the molecular target of DBM has not been discovered yet. DBM has good properties and no obvious side effects have been reported [35]. The site of action of DBM was suggested to be similar to ISRIB since it reduces ATF4 protein levels without affecting the phosphorylation of eIF2α. Given that DBM mimics the effects of ISRIB (i.e. partially restoring global protein synthesis levels and decreasing expression of ATF4 [35]), we speculate that the improvement observed in memory performance of our C9ORF72 model could be attributed to a similar molecular mechanism where protein synthesis restoration is essential to regulate synaptic plasticity and behavior [75].

Our proteomic analysis revealed that our G4C2 repeat mouse model recapitulates some proteomic alterations found in both ALS and FTD, and that DBM has important global effects in correcting the proteomic alterations observed in the disease model. Two main hits identified in AAV-66R mice treated with DBM were members of the Rab subfamily of small GTPases and the dDENN domain, identified as Rab2a and Gdi1. Rab2a, when bound to GTP, recruits and/or activates a variety of effector molecules controlling cellular trafficking, impacting processes such as autophagy [24, 89]. The autophagy pathway is a relevant node of the proteostasis network altered by C9ORF72 pathogenesis [41, 80] with important roles in synaptic function [39]. Gdi1 regulates the GDP-GTP exchange reaction of members of the Rab family, thus regulating vesicular trafficking [28]. Interestingly, Gdi1 knockout animals develop cognitive impairment associated with altered biogenesis and recycling of synaptic vesicles in the hippocampus [6] and its mutations are associated with X-linked general learning disability in humans [20]. Additionally, intermediate filaments vimentin (Vim) and alpha-internexin (Ina) were also impacted following DBM treatment in C9ORF72 mice. Poly(PR)-dipeptides were reported to bind mainly to intermediate filaments, thereby disassembling Vim [47]. Remarkably, Vim was shown to be protective under stress as it binds protein aggregates and granules formed by RNA-binding intrinsically disordered proteins and directs their asymmetric partitioning [62]. Additionally, degradation of vimentin bundles at perinucleus and dissociation of β-tubulin network is induced by poly(PR) treatment [72]. Importantly, the actin cytoskeleton has been extensively reported to regulate synaptic plasticity and brain function through different mechanisms, highlighting the regulation of dendritic spine dynamics [43, 46, 92]. Interestingly, actin cytoskeleton dynamics have been shown to regulate the ISR, by controlling eIF2α dephosphorylation [12, 13]. Remarkably, trazodone, a compound with similar activity to DBM, has important neuroprotective effects associated with the regulation of proteins involved in synaptic function, and metabolism, consistent with our current results [2]. Overall, our proteomic analysis suggests different possible mechanisms that may explain the improvement in our behavior phenotypes following DBM treatment in our C9ORF72 mouse model. Future studies will aim to study the possible contribution of our proteomic hits to C9ORF72 pathogenesis. In summary, we have demonstrated that oral administration of DBM improves cognitive performance in a mouse model of C9ORF72 pathogenesis, associated with proteomic changes that may be relevant to ALS/FTD.

## Authorship contribution statement

Claudio Hetz: Writing – Funding acquisition, conceptualization, original draft, resources, review & editing, project administration.

Paulina Torres, Grant Kauwe, Daniela Becerra, José I. Astorga, Matías Fuentealba, Guillermo Diaz, Luis Gonzalez, Vicente Valenzuela, Cameron Wehrfritz, Samah Shah, Joanna Bons, Yani Y. Ngwala: Investigation, visualization, data analysis.

Leonard Petrucelli, Tara E. Tracy: provided materials and gave experimental advice. Birgit Schilling: proteomic analysis.

## Consent for publication

All authors have agreed to publish the manuscript in this version.

## Data availability

Raw data and complete MS data sets have been uploaded to the Center for Computational Mass Spectrometry, MassIVE, and can be downloaded using the following FTP link: ftp://MSV000097847@massive-ftp.ucsd.edu or via the MassIVE website: https://massive.ucsd.edu/ProteoSAFe/dataset.jsp?task=633286dfbf224712bb0b25f6ed540756 (MassIVE ID: MSV000097847; ProteomeXchange ID: PXD063826).

[Note to the reviewers: To access the data repository MassIVE (UCSD) for MS data, please use: Username: MSV000097847_reviewer; Password: winter].

## Ethics approval

The experimental procedures involving mouse lines were approved by the Institutional Review Board for Animal Care of the Faculty of Medicine of the University of Chile (approved protocol CBA #18214-FMUCH).

## Acknowledgments

We thank Javiera Ponce for animal care supervision and Claudia Sepúlveda and Susana Manriquez for lab managing and administration. We thank Giovanna Mallucci (Altos Lab) for technical advice and help in experimental design in addition to provide DBM for the studies. We acknowledge the New York Genome Center (NYGC) ALS Consortium for generating the RNA-seq data utilized in this study. The datasets were obtained from the NCBI Gene Expression Omnibus (GSE137810). We thank the patients and their families for donation of tissue samples, which made this research possible.

## Supplementary legends

**Figure S1.**
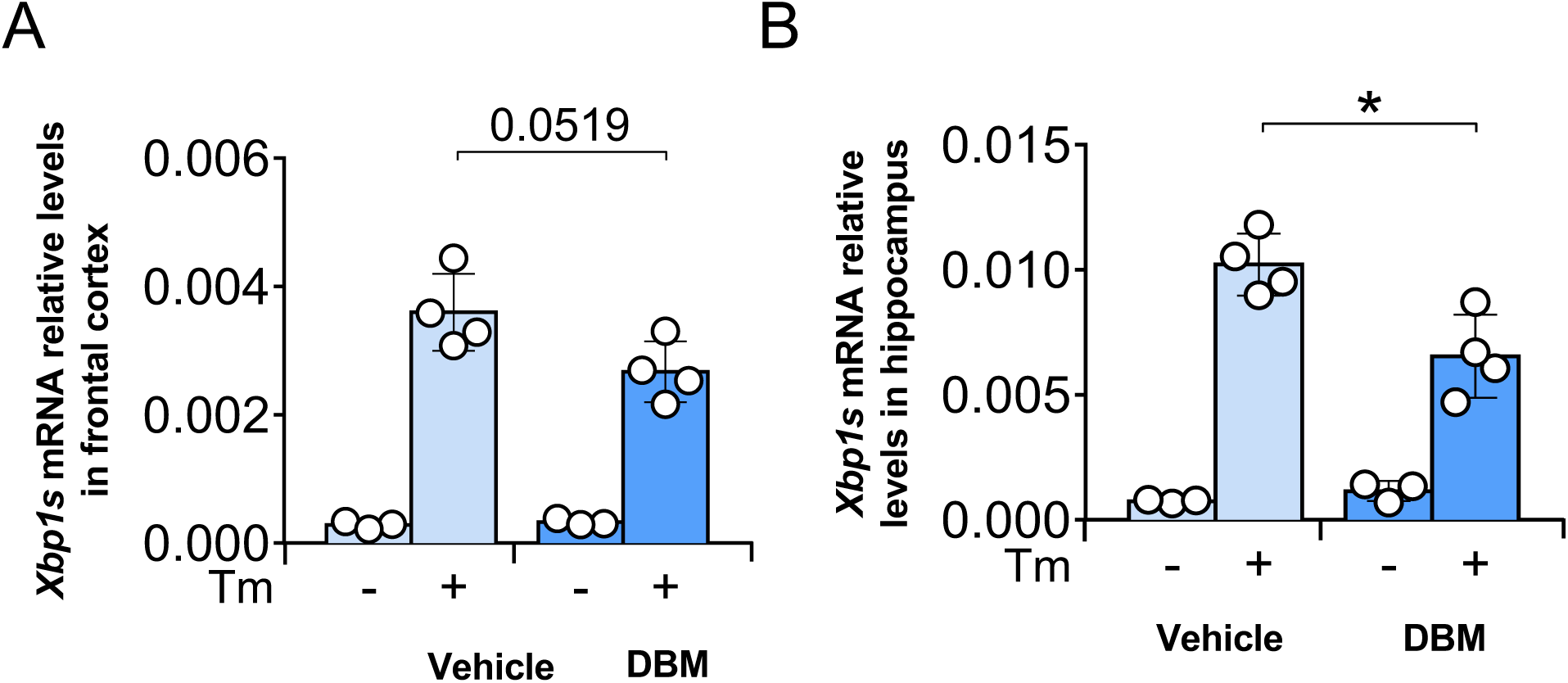
Hippocampal *Xbp1s* mRNA levels decreased after DBM treatment in mice under experimental ER stress. One-month-old mice were fed *ad libitum* with powdered chow containing 0.5% dibenzoylmethane (DBM) or vehicle for 7 days. Animals were then injected with 5 μg/g of tunicamycin (Tm), and brain tissue was collected 24 h later for biochemical analysis. *Xbp1s* mRNA levels were measured by qPCR in **(A)** frontal cortex and **(B)** hippocampus. All data were normalized with *Actin* mRNA levels. Each dot represents one animal. Data are presented as mean ± S.D. *, *p* < 0.05, one-way ANOVA followed by Tukey’s multiple comparisons test (n = 3-4).

**Figure S2.**
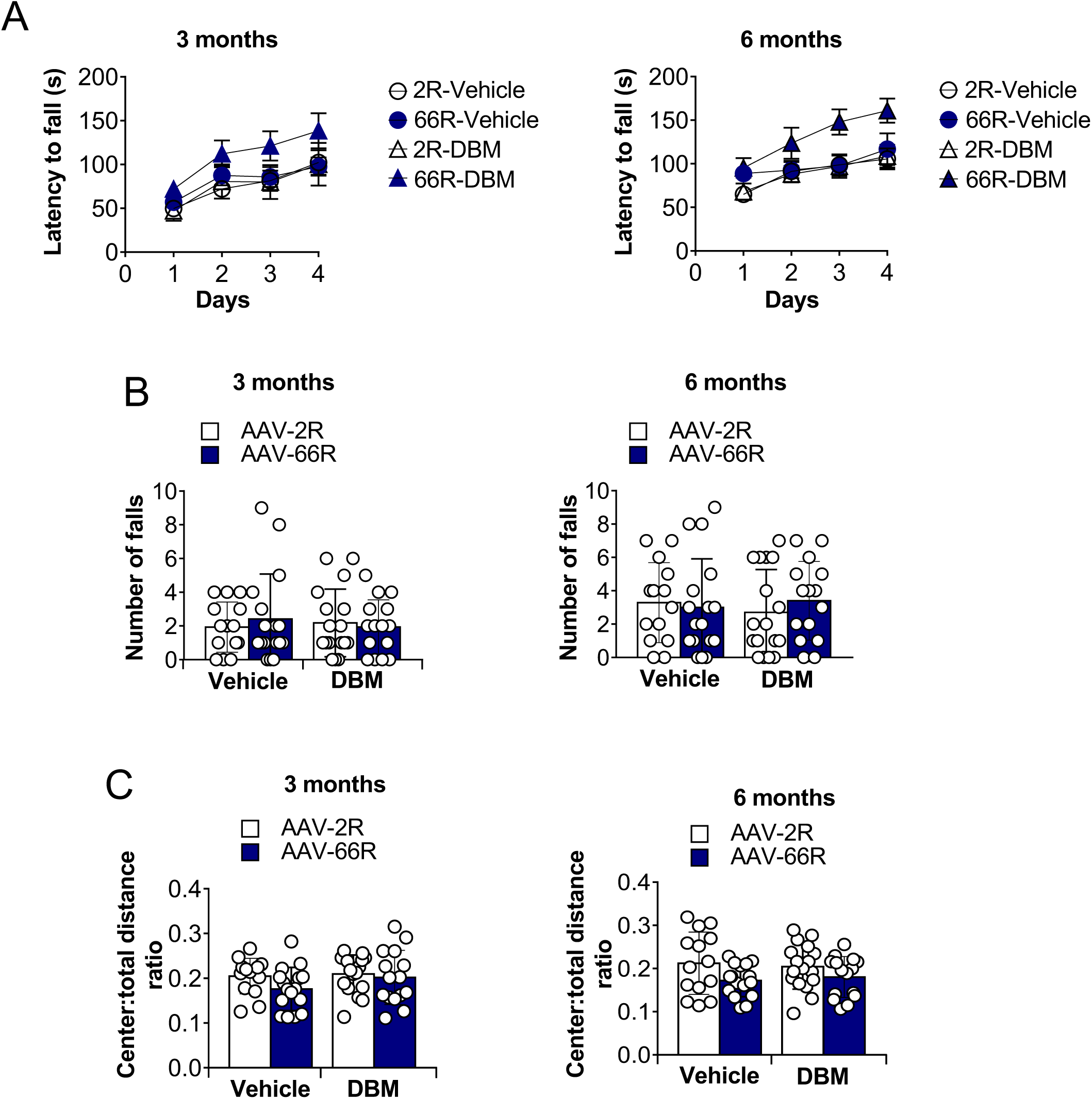
Behavioral testing in animals treated with DBM. Animals were injected with AAV-2R or AAV-66R at P0-1 and then fed with powdered chow containing 0.5% dibenzoylmethane (DBM) or vehicle *ad libitum* from the age of 1 month until the end of the experiment. **(A)** rotarod performance, **(B)** hanging wire test and **(C)** open field test, were performed at 3 and 6 months old. Each dot represents one animal. Data are presented as mean ± S.D. One-way ANOVA followed by Tukey’s multiple comparisons test (n = 14-17).

**Figure S3.**
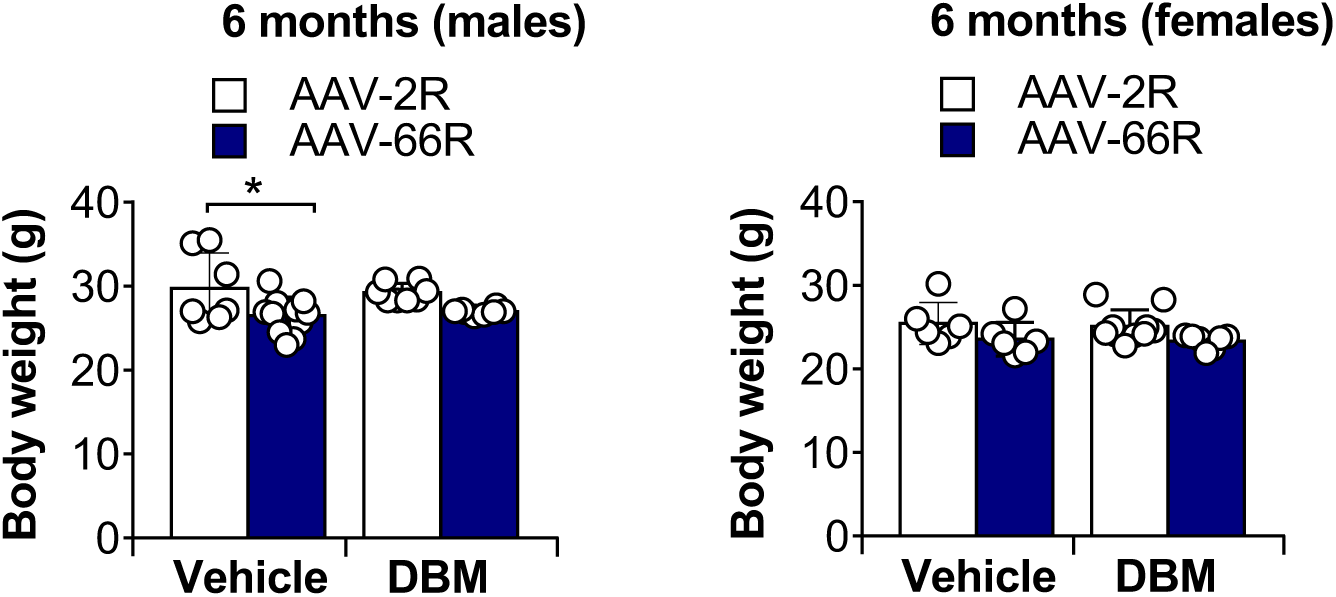
Body weight of C9orf72-FTD/ALS mice at 6-months of age. Animals injected with the AAV-2R or AAV-66R and then fed with powdered chow containing 0.5% DBM or vehicle *ad libitum* from the age of 1 month until the end of the experiment. Body weight was measured in animals at 6 months of age.

**Figure S4.**
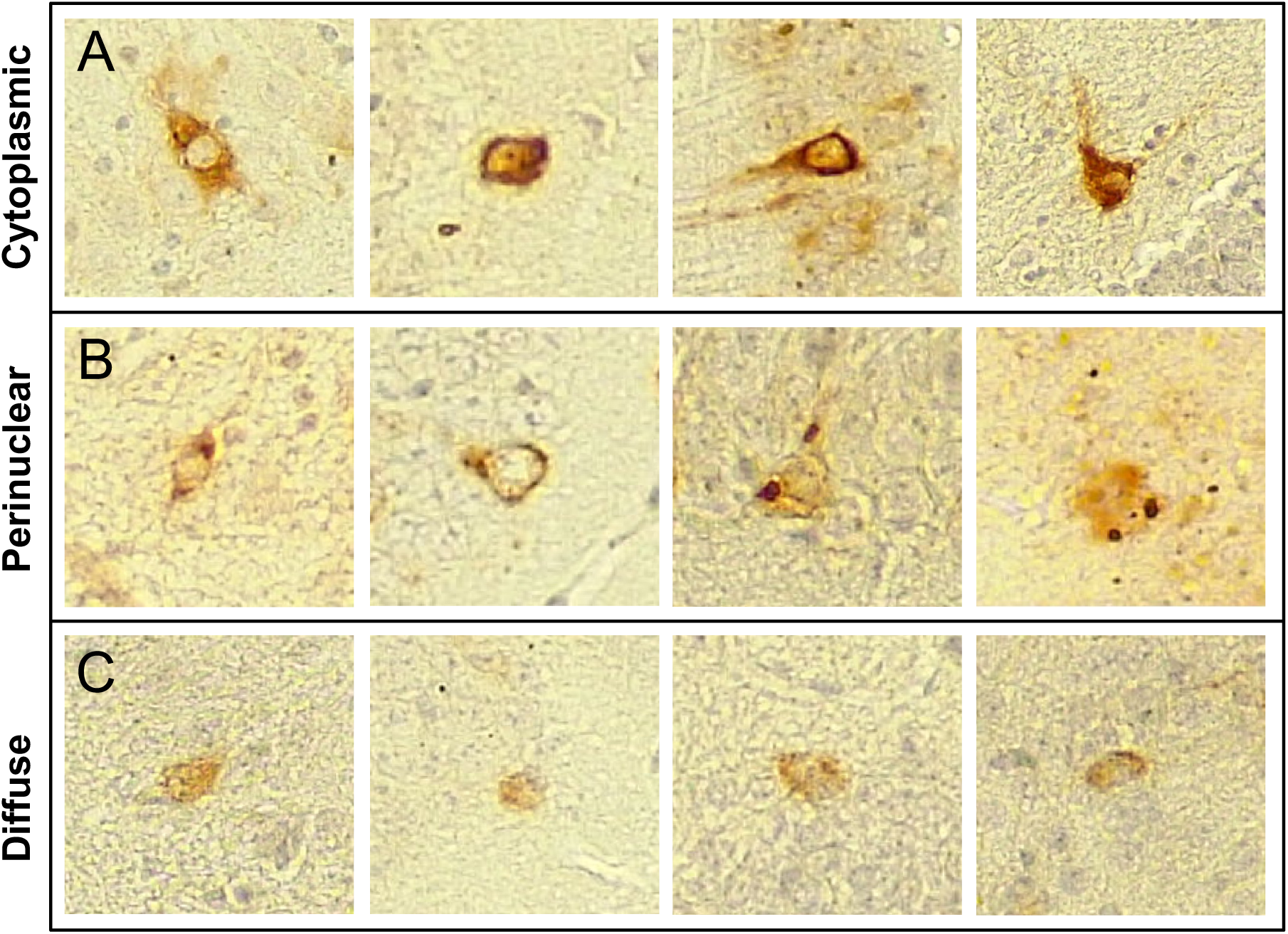
Representative images of inclusion classification. Brains from animals injected with AAV-66R were analyzed histologically to stain for poly(GA)-positive cells. Representative examples of **(A)** cytoplasmic, **(B)** perinuclear, and (**C)** diffuse distributions of poly(GA) inclusions in the are shown.

**Figure S5.**
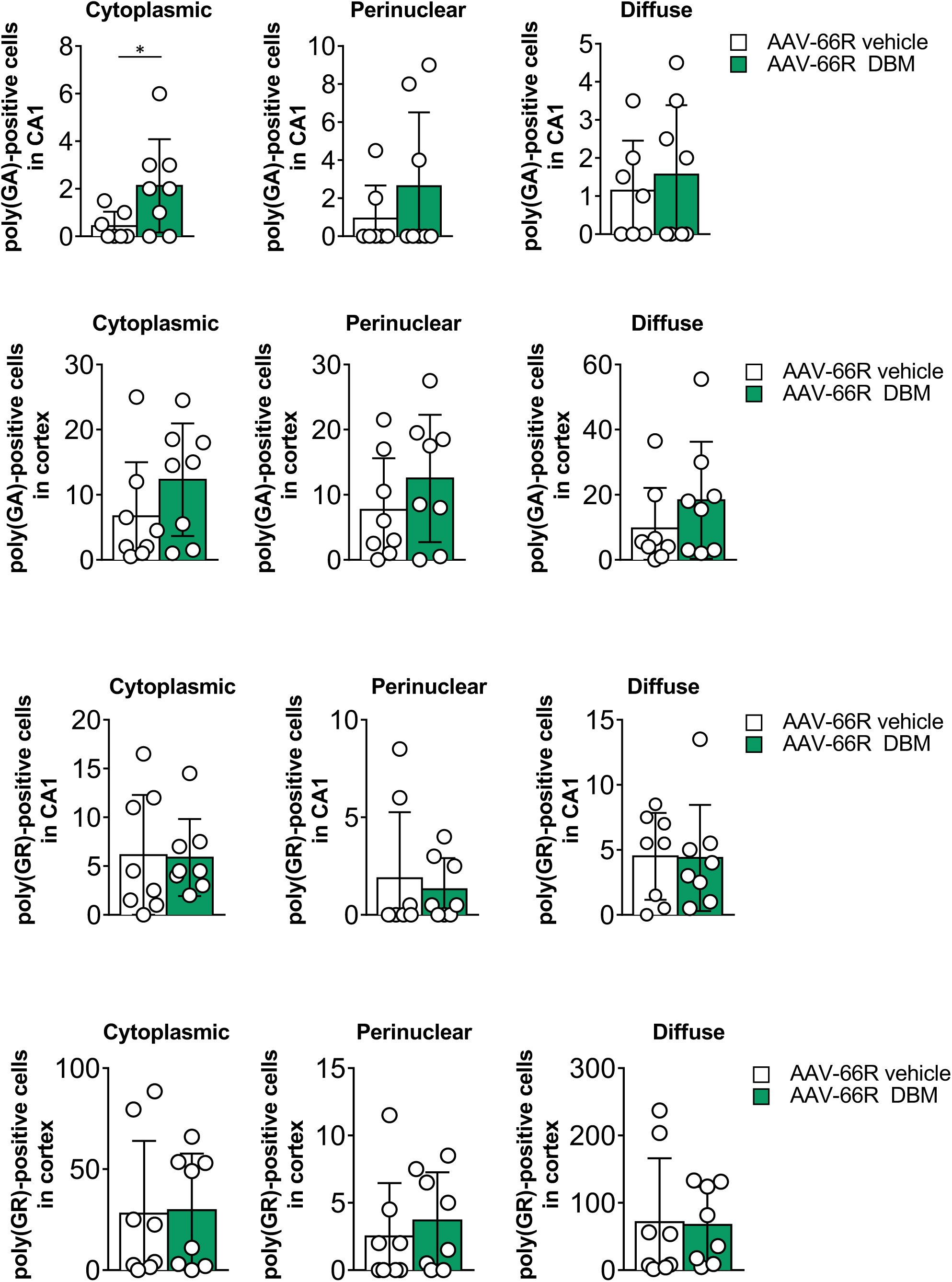
Cellular distribution quantification of poly(GA)-inclusions and poly(GR)-inclusions in CA1 and cortex. Animals were injected with AAV-2R or AAV-66R at P0-1 and then fed with powdered chow containing 0.5% DBM or vehicle *ad libitum* from the age of 1 month until the end of the experiment. Tissue was collected from animals at 6 months of age for histological analysis. Quantification of poly(GA)-and poly(GR)-positive cells in the cortex and hippocampus (CA1 region) of C9orf72-mediated FTD/ALS mice treated with vehicle or DBM. Each dot represents one animal (n = 8 per group). Data are presented as mean ± S.D. Unpaired-t test.

**Figure S6.**
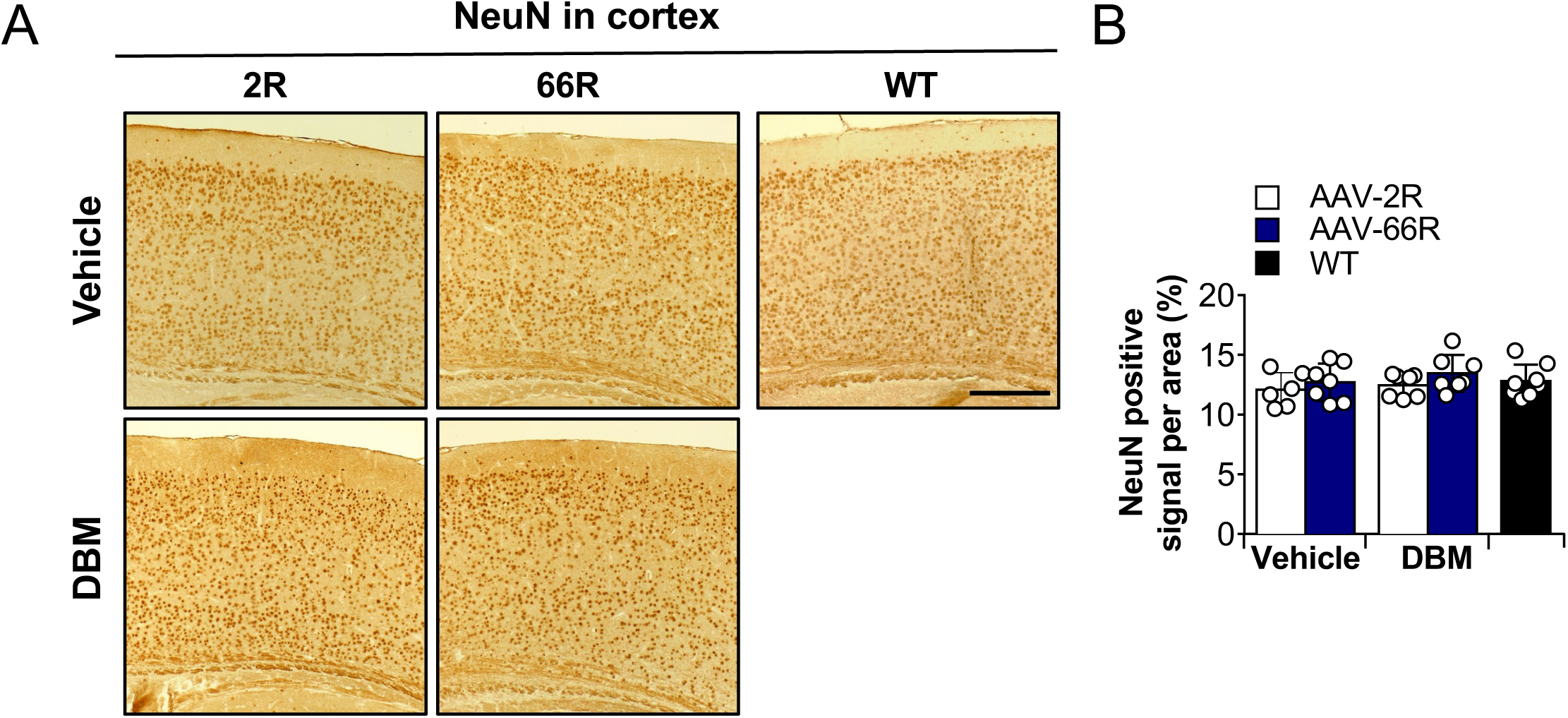
Neuronal survival in the brain cortex of AAV-66R mice. Animals were injected with the AAV-2R or AAV-66R at P0-1, and then fed with powdered chow containing 0.5% DBM or vehicle ad libitum from the age of 1 month until the end of the experiment. **(A)** At 6 months of age, brain tissue was collected for histological analysis. Representative images of NeuN-labeled cells in the brain cortex are presented. **(B)** Quantification of NeuN-positive cells in the cortex of C9ORF72-mediated FTD/ALS mice treated with vehicle or DBM. The percentage of NeuN-positive cells relative to the analyzed cortical area is shown. Each dot represents one animal (n = 8 per group). Scale bar (B): 300 µm.

**Figure S7.**
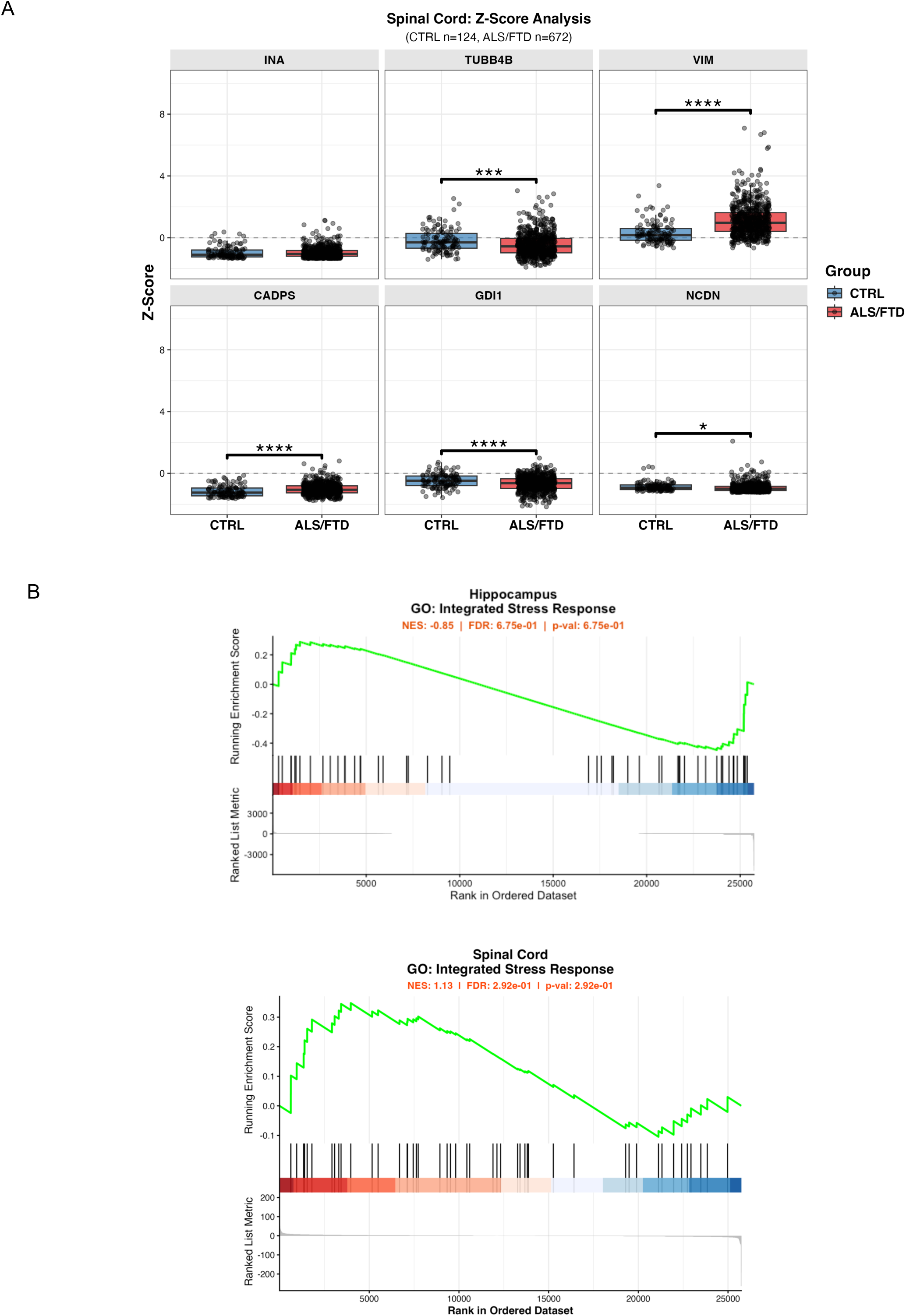
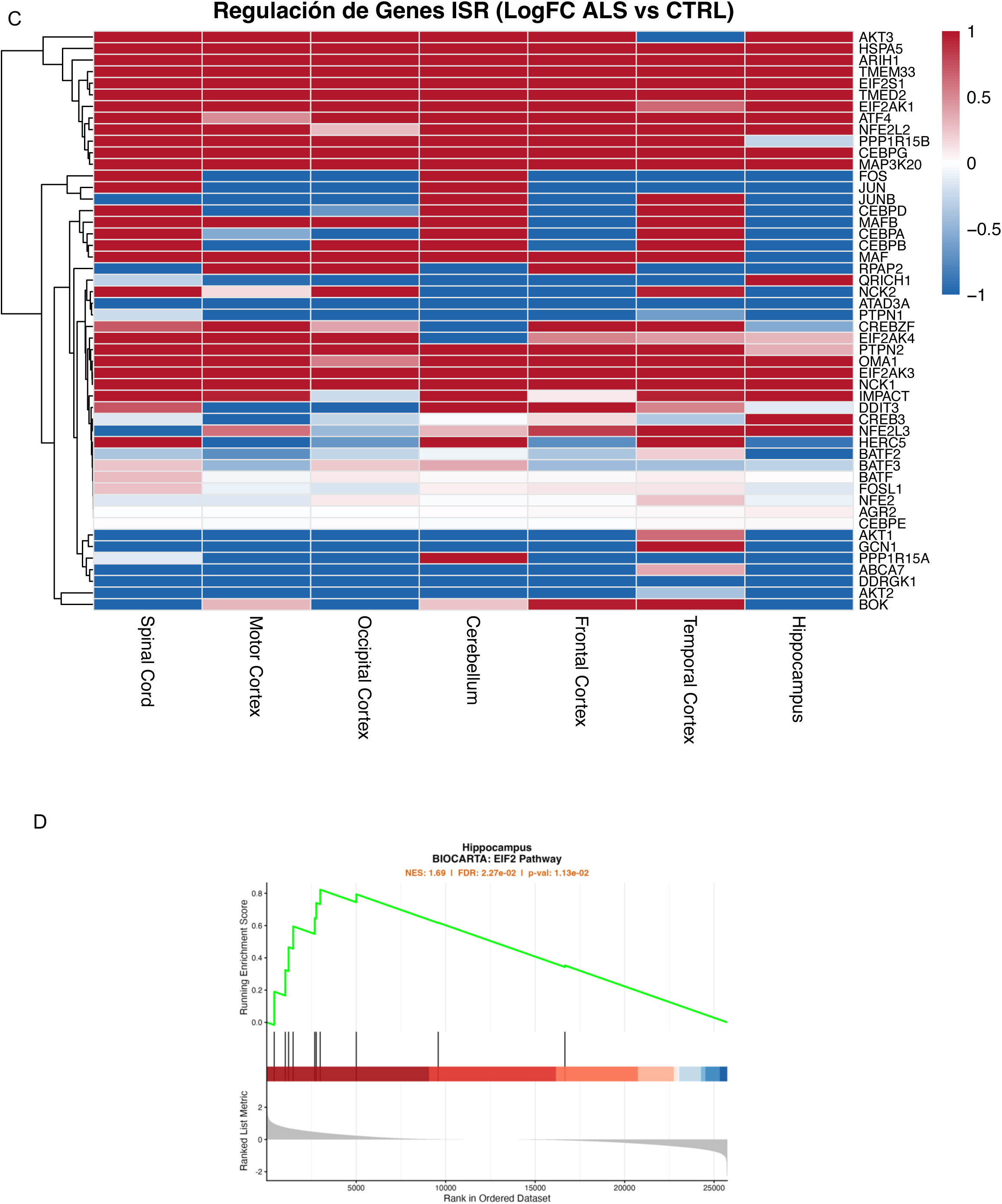
Analysis of human FTD/ALS gene expression databases. **(A)** Box-and-whisker plots depicting the differential expression (Z-score normalized) of DBM-corrected proteins (INA, TUBB4B, VIM, CADPS, GDI1, NCDN) in spinal cord tissue from control (CTRL, n = 124) and ALS/FTD (n = 672) subjects. The central line indicates the median, box limits represent the upper and lower quartiles, and whiskers extend to 1.5 times the interquartile range. Statistical significance was determined using an unpaired two-tailed t-test (* *p* < 0.05, *** *p* < 0.001, **** *p* < 0.0001). **(B)** Gene Set Enrichment Analysis (GSEA) plots for the Integrated Stress Response (ISR) biological process (GO:0140467) in the hippocampus and spinal cord. **(C)** Heatmap illustrating the Log2 Fold Change (LogFC, ALS vs. CTRL) of individual genes belonging to the ISR gene set (GO:0140467) across seven anatomical regions. Sample sizes for each region (CTRL / ALS-FTD) are: Spinal Cord (124/672), Motor Cortex (47/453), Occipital Cortex (11/63), Cerebellum (38/339), Frontal Cortex (55/372), Temporal Cortex (24/60), and Hippocampus (11/40). **(D)** GSEA plot showing significant positive enrichment of the eIF2 signaling pathway (Biocarta M6924, comprising 10 genes) in the hippocampus of ALS/FTD patients (NES = 1.69, FDR = 0.02).

---

**Table S1.**
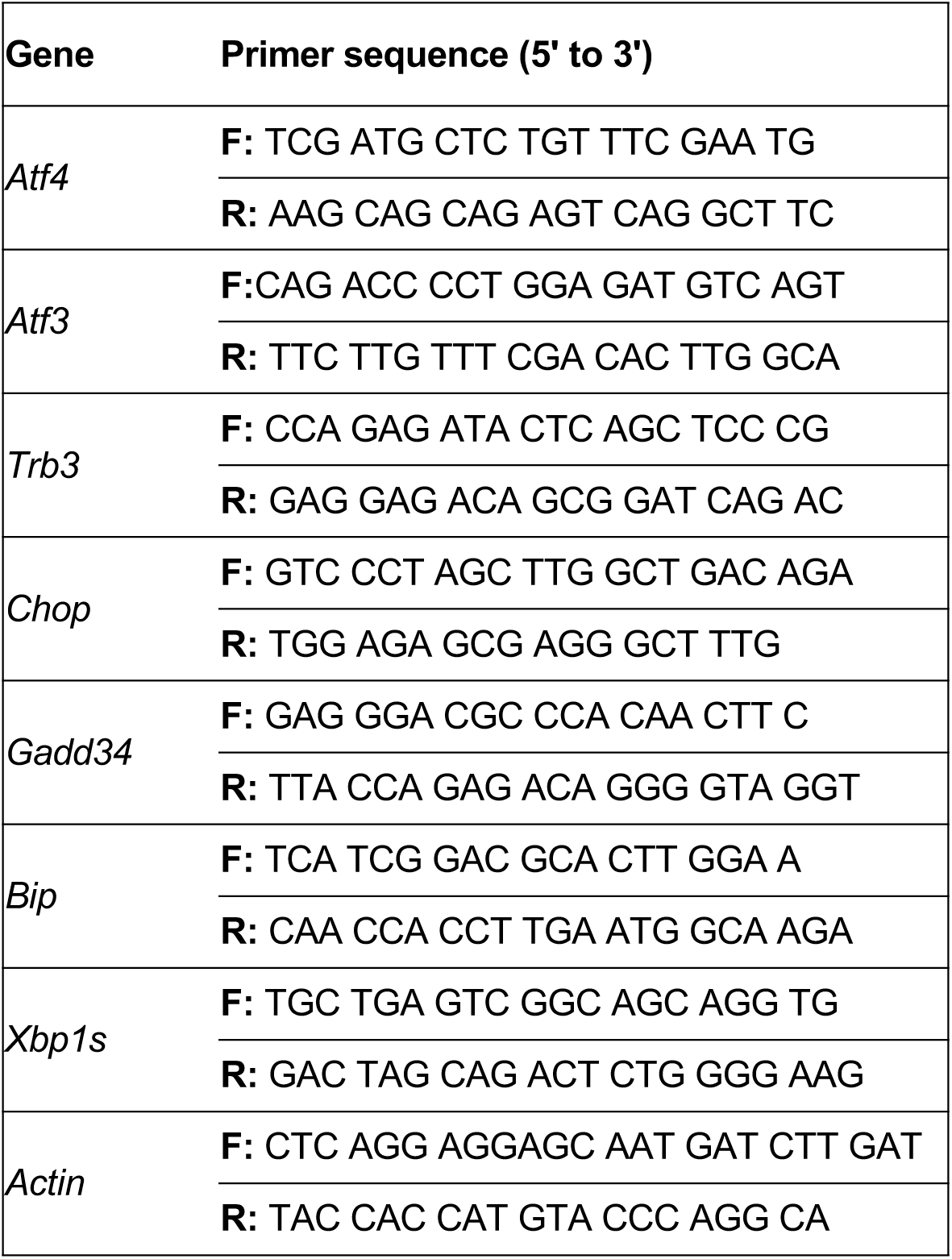
Primer used for RT-PCR. F: Forward; R: Reverse.

## References

1 Acosta-Alvear D, Harnoss JM, Walter P, Ashkenazi A (2025) Homeostasis control in health and disease by the unfolded protein response. Nat Rev Mol Cell Biol 26: 193–212 Doi 10.1038/s41580-024-00794-0

2 Albert-Gasco H, Smith HL, Alvarez-Castelao B, Swinden D, Halliday M, Janaki-Raman S, Butcher AJ, Mallucci GR (2024) Trazodone rescues dysregulated synaptic and mitochondrial nascent proteomes in prion neurodegeneration. Brain 147: 649–664 Doi 10.1093/brain/awad313

3 Anderson NS, Haynes CM (2020) Folding the Mitochondrial UPR into the Integrated Stress Response. Trends Cell Biol 30: 428–439 Doi 10.1016/j.tcb.2020.03.001

4 Ash PE, Bieniek KF, Gendron TF, Caulfield T, Lin WL, Dejesus-Hernandez M, van Blitterswijk MM, Jansen-West K, Paul JW, 3rd, Rademakers R et al (2013) Unconventional translation of C9ORF72 GGGGCC expansion generates insoluble polypeptides specific to c9FTD/ALS. Neuron 77: 639–646 Doi 10.1016/j.neuron.2013.02.004

5 Batra R, Lee CW (2017) Mouse Models of C9orf72 Hexanucleotide Repeat Expansion in Amyotrophic Lateral Sclerosis/ Frontotemporal Dementia. Front Cell Neurosci 11: 196 Doi 10.3389/fncel.2017.00196

6 Bianchi V, Farisello P, Baldelli P, Meskenaite V, Milanese M, Vecellio M, Muhlemann S, Lipp HP, Bonanno G, Benfenati F et al (2009) Cognitive impairment in Gdi1-deficient mice is associated with altered synaptic vesicle pools and short-term synaptic plasticity, and can be corrected by appropriate learning training. Hum Mol Genet 18: 105–117 Doi 10.1093/hmg/ddn321

7 Bravo-Jimenez MA, Sharma S, Karimi-Abdolrezaee S (2025) The integrated stress response in neurodegenerative diseases. Mol Neurodegener 20: 20 Doi 10.1186/s13024-025-00811-6

8 Burberry A, Wells MF, Limone F, Couto A, Smith KS, Keaney J, Gillet G, van Gastel N, Wang JY, Pietilainen O et al (2020) C9orf72 suppresses systemic and neural inflammation induced by gut bacteria. Nature 582: 89–94 Doi 10.1038/s41586-020-2288-7

9 Burger T (2018) Gentle Introduction to the Statistical Foundations of False Discovery Rate in Quantitative Proteomics. Journal of proteome research 17: 12–22 Doi 10.1021/acs.jproteome.7b00170

10 Burrell JR, Halliday GM, Kril JJ, Ittner LM, Gotz J, Kiernan MC, Hodges JR (2016) The frontotemporal dementia-motor neuron disease continuum. Lancet 388: 919–931 Doi 10.1016/S0140-6736(16)00737-6

11 Cabral-Miranda F, Tamburini G, Martinez G, Ardiles AO, Medinas DB, Gerakis Y, Hung MD, Vidal R, Fuentealba M, Miedema T et al (2022) Unfolded protein response IRE1/XBP1 signaling is required for healthy mammalian brain aging. EMBO J 41: e111952 Doi 10.15252/embj.2022111952

12 Chambers JE, Dalton LE, Clarke HJ, Malzer E, Dominicus CS, Patel V, Moorhead G, Ron D, Marciniak SJ (2015) Actin dynamics tune the integrated stress response by regulating eukaryotic initiation factor 2alpha dephosphorylation. eLife 4: Doi 10.7554/eLife.04872

13 Chen R, Rato C, Yan Y, Crespillo-Casado A, Clarke HJ, Harding HP, Marciniak SJ, Read RJ, Ron D (2015) G-actin provides substrate-specificity to eukaryotic initiation factor 2alpha holophosphatases. eLife 4: Doi 10.7554/eLife.04871

14 Chen-Plotkin AS, Lee VM, Trojanowski JQ (2010) TAR DNA-binding protein 43 in neurodegenerative disease. Nat Rev Neurol 6: 211–220 Doi 10.1038/nrneurol.2010.18

15 Cheng W, Wang S, Mestre AA, Fu C, Makarem A, Xian F, Hayes LR, Lopez-Gonzalez R, Drenner K, Jiang J et al (2018) C9ORF72 GGGGCC repeat-associated non-AUG translation is upregulated by stress through eIF2alpha phosphorylation. Nat Commun 9: 51 Doi 10.1038/s41467-017-02495-z

16 Chew J, Gendron TF, Prudencio M, Sasaguri H, Zhang YJ, Castanedes-Casey M, Lee CW, Jansen-West K, Kurti A, Murray ME et al (2015) Neurodegeneration. C9ORF72 repeat expansions in mice cause TDP-43 pathology, neuronal loss, and behavioral deficits. Science 348: 1151–1154 Doi 10.1126/science.aaa9344

17 Chou A, Krukowski K, Jopson T, Zhu PJ, Costa-Mattioli M, Walter P, Rosi S (2017) Inhibition of the integrated stress response reverses cognitive deficits after traumatic brain injury. Proc Natl Acad Sci U S A 114: E6420–E6426 Doi 10.1073/pnas.1707661114

18 Collins BC, Hunter CL, Liu Y, Schilling B, Rosenberger G, Bader SL, Chan DW, Gibson BW, Gingras AC, Held JM et al (2017) Multi-laboratory assessment of reproducibility, qualitative and quantitative performance of SWATH-mass spectrometry. Nature communications 8: 291 Doi 10.1038/s41467-017-00249-5

19 Costa-Mattioli M, Walter P (2020) The integrated stress response: From mechanism to disease. Science 368: Doi 10.1126/science.aat5314

20 D’Adamo P, Menegon A, Lo Nigro C, Grasso M, Gulisano M, Tamanini F, Bienvenu T, Gedeon AK, Oostra B, Wu SK et al (1998) Mutations in GDI1 are responsible for X-linked non-specific mental retardation. Nat Genet 19: 134–139 Doi 10.1038/487

21 Dafinca R, Scaber J, Ababneh N, Lalic T, Weir G, Christian H, Vowles J, Douglas AG, Fletcher-Jones A, Browne C et al (2016) C9orf72 Hexanucleotide Expansions Are Associated with Altered Endoplasmic Reticulum Calcium Homeostasis and Stress Granule Formation in Induced Pluripotent Stem Cell-Derived Neurons from Patients with Amyotrophic Lateral Sclerosis and Frontotemporal Dementia. Stem Cells 34: 2063–2078 Doi 10.1002/stem.2388

22 DeJesus-Hernandez M, Mackenzie IR, Boeve BF, Boxer AL, Baker M, Rutherford NJ, Nicholson AM, Finch NA, Flynn H, Adamson J et al (2011) Expanded GGGGCC hexanucleotide repeat in noncoding region of C9ORF72 causes chromosome 9p-linked FTD and ALS. Neuron 72: 245–256 Doi 10.1016/j.neuron.2011.09.011

23 Dhir N, Jain A, Sharma AR, Prakash A, Radotra BD, Medhi B (2023) PERK inhibitor, GSK2606414, ameliorates neuropathological damage, memory and motor functional impairments in cerebral ischemia via PERK/p-eIF2a/ATF4/CHOP signaling. Metab Brain Dis 38: 1177–1192 Doi 10.1007/s11011-023-01183-w

24 Ding X, Jiang X, Tian R, Zhao P, Li L, Wang X, Chen S, Zhu Y, Mei M, Bao S et al (2019) RAB2 regulates the formation of autophagosome and autolysosome in mammalian cells. Autophagy 15: 1774–1786 Doi 10.1080/15548627.2019.1596478

25 Elliott E, Bailey O, Waldron FM, Hardingham GE, Chandran S, Gregory JM (2020) Therapeutic Targeting of Proteostasis in Amyotrophic Lateral Sclerosis-a Systematic Review and Meta-Analysis of Preclinical Research. Front Neurosci 14: 511 Doi 10.3389/fnins.2020.00511

26 Espina M, Di Franco N, Branas-Navarro M, Navarro IR, Brito V, Lopez-Molina L, Costas-Insua C, Guzman M, Gines S (2023) The GRP78-PERK axis contributes to memory and synaptic impairments in Huntington’s disease R6/1 mice. Neurobiol Dis 184: 106225 Doi 10.1016/j.nbd.2023.106225

27 Gami-Patel P, van Dijken I, Meeter LH, Melhem S, Morrema THJ, Scheper W, van Swieten JC, Rozemuller AJM, Dijkstra AA, Hoozemans JJM (2021) Unfolded protein response activation in C9orf72 frontotemporal dementia is associated with dipeptide pathology and granulovacuolar degeneration in granule cells. Brain Pathol 31: 163–173 Doi 10.1111/bpa.12894

28 Garrett MD, Zahner JE, Cheney CM, Novick PJ (1994) GDI1 encodes a GDP dissociation inhibitor that plays an essential role in the yeast secretory pathway. EMBO J 13: 1718–1728 Doi 10.1002/j.1460-2075.1994.tb06436.x

29 Gendron TF, Belzil VV, Zhang YJ, Petrucelli L (2014) Mechanisms of toxicity in C9FTLD/ALS. Acta Neuropathol 127: 359–376 Doi 10.1007/s00401-013-1237-z

30 Gillet LC, Navarro P, Tate S, Rost H, Selevsek N, Reiter L, Bonner R, Aebersold R (2012) Targeted data extraction of the MS/MS spectra generated by data-independent acquisition: a new concept for consistent and accurate proteome analysis. Molecular & cellular proteomics: MCP 11: O111 016717 Doi 10.1074/mcp.O111.016717

31 Gotoh S, Mori K, Fujino Y, Kawabe Y, Yamashita T, Omi T, Nagata K, Tagami S, Nagai Y, Ikeda M (2024) eIF5 stimulates the CUG initiation of RAN translation of poly-GA dipeptide repeat protein (DPR) in C9orf72 FTLD/ALS. J Biol Chem 300: 105703 Doi 10.1016/j.jbc.2024.105703

32 Graham GJ, Rennie JS (1988) A cross-sectional study of the effects of alcohol on the protein profile of rat lingual epithelium. Arch Oral Biol 33: 631–634 Doi 10.1016/0003-9969(88)90115-x

33 Green KM, Glineburg MR, Kearse MG, Flores BN, Linsalata AE, Fedak SJ, Goldstrohm AC, Barmada SJ, Todd PK (2017) RAN translation at C9orf72-associated repeat expansions is selectively enhanced by the integrated stress response. Nat Commun 8: 2005 Doi 10.1038/s41467-017-02200-0

34 Halliday M, Radford H, Sekine Y, Moreno J, Verity N, le Quesne J, Ortori CA, Barrett DA, Fromont C, Fischer PM et al (2015) Partial restoration of protein synthesis rates by the small molecule ISRIB prevents neurodegeneration without pancreatic toxicity. Cell Death Dis 6: e1672 Doi 10.1038/cddis.2015.49

35 Halliday M, Radford H, Zents KAM, Molloy C, Moreno JA, Verity NC, Smith E, Ortori CA, Barrett DA, Bushell M et al (2017) Repurposed drugs targeting eIF2alpha-P-mediated translational repression prevent neurodegeneration in mice. Brain 140: 1768–1783 Doi 10.1093/brain/awx074

36 Hardiman O, Al-Chalabi A, Chio A, Corr EM, Logroscino G, Robberecht W, Shaw PJ, Simmons Z, van den Berg LH (2017) Amyotrophic lateral sclerosis. Nature reviews Disease primers 3: 17071 Doi 10.1038/nrdp.2017.71

37 Harper NS, Sharpe JL, Speranza J, Gulia R, Chen JX, Allen SP, Atwal MS, Pickering-Brown S, Livesey MR, Bennett CL et al (2026) Targeting the integrated stress response or Ataxin-2 alleviates neurodegeneration in PolyGR models of C9orf72 associated frontotemporal dementia and amyotrophic lateral sclerosis. Acta neuropathologica communications 14: Doi 10.1186/s40478-026-02301-2

38 Hartmann H, Hornburg D, Czuppa M, Bader J, Michaelsen M, Farny D, Arzberger T, Mann M, Meissner F, Edbauer D (2018) Proteomics and C9orf72 neuropathology identify ribosomes as poly-GR/PR interactors driving toxicity. Life Sci Alliance 1: e201800070 Doi 10.26508/lsa.201800070

39 Hetz C (2021) Adapting the proteostasis capacity to sustain brain healthspan. Cell 184: 1545–1560 Doi 10.1016/j.cell.2021.02.007

40 Hetz C, Axten JM, Patterson JB (2019) Pharmacological targeting of the unfolded protein response for disease intervention. Nat Chem Biol 15: 764–775 Doi 10.1038/s41589-019-0326-2

41 Houghton OH, Mizielinska S, Gomez-Suaga P (2022) The Interplay Between Autophagy and RNA Homeostasis: Implications for Amyotrophic Lateral Sclerosis and Frontotemporal Dementia. Front Cell Dev Biol 10: 838402 Doi 10.3389/fcell.2022.838402

42 Kanekura K, Yagi T, Cammack AJ, Mahadevan J, Kuroda M, Harms MB, Miller TM, Urano F (2016) Poly-dipeptides encoded by the C9ORF72 repeats block global protein translation. Hum Mol Genet 25: 1803–1813 Doi 10.1093/hmg/ddw052

43 Konietzny A, Bar J, Mikhaylova M (2017) Dendritic Actin Cytoskeleton: Structure, Functions, and Regulations. Front Cell Neurosci 11: 147 Doi 10.3389/fncel.2017.00147

44 Kramer NJ, Haney MS, Morgens DW, Jovicic A, Couthouis J, Li A, Ousey J, Ma R, Bieri G, Tsui CK et al (2018) CRISPR-Cas9 screens in human cells and primary neurons identify modifiers of C9ORF72 dipeptide-repeat-protein toxicity. Nat Genet 50: 603–612 Doi 10.1038/s41588-018-0070-7

45 Kriachkov V, Ormsby AR, Kusnadi EP, McWilliam HEG, Mintern JD, Amarasinghe SL, Ritchie ME, Furic L, Hatters DM (2023) Arginine-rich C9ORF72 ALS proteins stall ribosomes in a manner distinct from a canonical ribosome-associated quality control substrate. J Biol Chem 299: 102774 Doi 10.1016/j.jbc.2022.102774

46 Lei W, Omotade OF, Myers KR, Zheng JQ (2016) Actin cytoskeleton in dendritic spine development and plasticity. Curr Opin Neurobiol 39: 86–92 Doi 10.1016/j.conb.2016.04.010

47 Lin Y, Mori E, Kato M, Xiang S, Wu L, Kwon I, McKnight SL (2016) Toxic PR Poly-Dipeptides Encoded by the C9orf72 Repeat Expansion Target LC Domain Polymers. Cell 167: 789–802 e712 Doi 10.1016/j.cell.2016.10.003

48 Ling SC, Polymenidou M, Cleveland DW (2013) Converging mechanisms in ALS and FTD: disrupted RNA and protein homeostasis. Neuron 79: 416–438 Doi 10.1016/j.neuron.2013.07.033

49 Liu F, Morderer D, Wren MC, Vettleson-Trutza SA, Wang Y, Rabichow BE, Salemi MR, Phinney BS, Oskarsson B, Dickson DW et al (2022) Proximity proteomics of C9orf72 dipeptide repeat proteins identifies molecular chaperones as modifiers of poly-GA aggregation. Acta Neuropathol Commun 10: 22 Doi 10.1186/s40478-022-01322-x

50 Loveland AB, Svidritskiy E, Susorov D, Lee S, Park A, Zvornicanin S, Demo G, Gao FB, Korostelev AA (2022) Ribosome inhibition by C9ORF72-ALS/FTD-associated poly-PR and poly-GR proteins revealed by cryo-EM. Nat Commun 13: 2776 Doi 10.1038/s41467-022-30418-0

51 Mercado G, Castillo V, Soto P, Lopez N, Axten JM, Sardi SP, Hoozemans JJM, Hetz C (2018) Targeting PERK signaling with the small molecule GSK2606414 prevents neurodegeneration in a model of Parkinson’s disease. Neurobiology of disease 112: 136–148 Doi 10.1016/j.nbd.2018.01.004

52 Mizielinska S, Hautbergue GM, Gendron TF, van Blitterswijk M, Hardiman O, Ravits J, Isaacs AM, Rademakers R (2025) Amyotrophic lateral sclerosis caused by hexanucleotide repeat expansions in C9orf72: from genetics to therapeutics. Lancet Neurol 24: 261–274 Doi 10.1016/S1474-4422(25)00026-2

53 Moens TG, Niccoli T, Wilson KM, Atilano ML, Birsa N, Gittings LM, Holbling BV, Dyson MC, Thoeng A, Neeves J et al (2019) C9orf72 arginine-rich dipeptide proteins interact with ribosomal proteins in vivo to induce a toxic translational arrest that is rescued by eIF1A. Acta Neuropathol 137: 487–500 Doi 10.1007/s00401-018-1946-4

54 Mordes DA, Morrison BM, Ament XH, Cantrell C, Mok J, Eggan P, Xue C, Wang JY, Eggan K, Rothstein JD (2020) Absence of Survival and Motor Deficits in 500 Repeat C9ORF72 BAC Mice. Neuron 108: 775–783 e774 Doi 10.1016/j.neuron.2020.08.009

55 Moreno JA, Halliday M, Molloy C, Radford H, Verity N, Axten JM, Ortori CA, Willis AE, Fischer PM, Barrett DA et al (2013) Oral treatment targeting the unfolded protein response prevents neurodegeneration and clinical disease in prion-infected mice. Sci Transl Med 5: 206ra138 Doi 10.1126/scitranslmed.3006767

56 Moreno JA, Radford H, Peretti D, Steinert JR, Verity N, Martin MG, Halliday M, Morgan J, Dinsdale D, Ortori CA et al (2012) Sustained translational repression by eIF2alpha-P mediates prion neurodegeneration. Nature 485: 507–511 Doi 10.1038/nature11058

57 Moreno-Jimenez L, Benito-Martin MS, Sanclemente-Alaman I, Matias-Guiu JA, Sancho-Bielsa F, Canales-Aguirre A, Mateos-Diaz JC, Matias-Guiu J, Aguilar J, Gomez-Pinedo U (2024) Murine experimental models of amyotrophic lateral sclerosis: an update. Neurologia (Engl Ed) 39: 282–291 Doi 10.1016/j.nrleng.2021.07.004

58 Oliveira MM, Lourenco MV, Longo F, Kasica NP, Yang W, Ureta G, Ferreira DDP, Mendonca PHJ, Bernales S, Ma Tet al (2021) Correction of eIF2-dependent defects in brain protein synthesis, synaptic plasticity, and memory in mouse models of Alzheimer’s disease. Sci Signal 14: Doi 10.1126/scisignal.abc5429

59 Parakh S, Atkin JD (2016) Protein folding alterations in amyotrophic lateral sclerosis. Brain Res 1648: 633–649 Doi 10.1016/j.brainres.2016.04.010

60 Parameswaran J, Zhang N, Braems E, Tilahun K, Pant DC, Yin K, Asress S, Heeren K, Banerjee A, Davis E et al (2023) Antisense, but not sense, repeat expanded RNAs activate PKR/eIF2alpha-dependent ISR in C9ORF72 FTD/ALS. Elife 12: Doi 10.7554/eLife.85902

61 Pareja-Navarro KA, King CD, Kauwe G, Ngwala YY, Lokitiyakul D, Wong I, Vira A, Liu Y, Chen JH, Sharma M et al (2026) Tau oligomers modulate synapse fate by eliciting progressive bipartite synapse dysregulation and synapse loss. Molecular neurodegeneration 21: 13 Doi 10.1186/s13024-026-00928-2

62 Pattabiraman S, Azad GK, Amen T, Brielle S, Park JE, Sze SK, Meshorer E, Kaganovich D (2020) Vimentin protects differentiating stem cells from stress. Sci Rep 10: 19525 Doi 10.1038/s41598-020-76076-4

63 Peters OM, Ghasemi M, Brown RH, Jr. (2015) Emerging mechanisms of molecular pathology in ALS. J Clin Invest 125: 1767–1779 Doi 10.1172/JCI71601

64 Radford H, Moreno JA, Verity N, Halliday M, Mallucci GR (2015) PERK inhibition prevents tau-mediated neurodegeneration in a mouse model of frontotemporal dementia. Acta Neuropathol 130: 633–642 Doi 10.1007/s00401-015-1487-z

65 Renton AE, Majounie E, Waite A, Simon-Sanchez J, Rollinson S, Gibbs JR, Schymick JC, Laaksovirta H, van Swieten JC, Myllykangas L et al (2011) A hexanucleotide repeat expansion in C9ORF72 is the cause of chromosome 9p21-linked ALS-FTD. Neuron 72: 257–268 Doi 10.1016/j.neuron.2011.09.010

66 Sahana TG, Chase KJ, Liu F, Lloyd TE, Rossoll W, Zhang K (2023) c-Jun N-Terminal Kinase Promotes Stress Granule Assembly and Neurodegeneration in C9orf72-Mediated ALS and FTD. J Neurosci 43: 3186–3197 Doi 10.1523/JNEUROSCI.1799-22.2023

67 Saxena S, Cabuy E, Caroni P (2009) A role for motoneuron subtype-selective ER stress in disease manifestations of FALS mice. Nature neuroscience 12: 627–636 Doi 10.1038/nn.2297

68 Schilling B, Gibson BW, Hunter CL (2017) Generation of High-Quality SWATH((R)) Acquisition Data for Label-free Quantitative Proteomics Studies Using TripleTOF((R)) Mass Spectrometers. Methods Mol Biol 1550: 223–233 Doi 10.1007/978-1-4939-6747-6_16

69 Sekine Y, Zyryanova A, Crespillo-Casado A, Fischer PM, Harding HP, Ron D (2015) Stress responses. Mutations in a translation initiation factor identify the target of a memory-enhancing compound. Science 348: 1027–1030 Doi 10.1126/science.aaa6986

70 Sellier C, Campanari ML, Julie Corbier C, Gaucherot A, Kolb-Cheynel I, Oulad-Abdelghani M, Ruffenach F, Page A, Ciura S, Kabashi E et al (2016) Loss of C9ORF72 impairs autophagy and synergizes with polyQ Ataxin-2 to induce motor neuron dysfunction and cell death. EMBO J 35: 1276–1297 Doi 10.15252/embj.201593350

71 Shahheydari H, Ragagnin A, Walker AK, Toth RP, Vidal M, Jagaraj CJ, Perri ER, Konopka A, Sultana JM, Atkin JD (2017) Protein Quality Control and the Amyotrophic Lateral Sclerosis/Frontotemporal Dementia Continuum. Front Mol Neurosci 10: 119 Doi 10.3389/fnmol.2017.00119

72 Shiota T, Nagata R, Kikuchi S, Nanaura H, Matsubayashi M, Nakanishi M, Kobashigawa S, Isozumi N, Kiriyama T, Nagayama K et al (2022) C9orf72-Derived Proline:Arginine Poly-Dipeptides Modulate Cytoskeleton and Mechanical Stress Response. Front Cell Dev Biol 10: 750829 Doi 10.3389/fcell.2022.750829

73 Sidrauski C, Acosta-Alvear D, Khoutorsky A, Vedantham P, Hearn BR, Li H, Gamache K, Gallagher CM, Ang KK, Wilson C et al (2013) Pharmacological brake-release of mRNA translation enhances cognitive memory. eLife 2: e00498 Doi 10.7554/eLife.00498

74 Sonobe Y, Ghadge G, Masaki K, Sendoel A, Fuchs E, Roos RP (2018) Translation of dipeptide repeat proteins from the C9ORF72 expanded repeat is associated with cellular stress. Neurobiol Dis 116: 155–165 Doi 10.1016/j.nbd.2018.05.009

75 Sossin WS, Costa-Mattioli M (2019) Translational Control in the Brain in Health and Disease. Cold Spring Harbor perspectives in biology 11: Doi 10.1101/cshperspect.a032912

76 Storey JD (2002) A Direct Approach to False Discovery Rates. Journal of the Royal Statistical Society Series B: Statistical Methodology 64: 479–498 Doi 10.1111/1467-9868.00346

77 Sultana J, Ragagnin AMG, Parakh S, Saravanabavan S, Soo KY, Vidal M, Jagaraj CJ, Ding K, Wu S, Shadfar S et al (2024) C9orf72-Associated Dipeptide Repeat Expansions Perturb ER-Golgi Vesicular Trafficking, Inducing Golgi Fragmentation and ER Stress, in ALS/FTD. Molecular neurobiology 61: 10318–10338 Doi 10.1007/s12035-024-04187-4

78 Tao Z, Wang H, Xia Q, Li K, Li K, Jiang X, Xu G, Wang G, Ying Z (2015) Nucleolar stress and impaired stress granule formation contribute to C9orf72 RAN translation-induced cytotoxicity. Hum Mol Genet 24: 2426–2441 Doi 10.1093/hmg/ddv005

79 Taylor JP, Brown RH, Jr., Cleveland DW (2016) Decoding ALS: from genes to mechanism. Nature 539: 197–206 Doi 10.1038/nature20413

80 Torres P, Cabral-Miranda F, Gonzalez-Teuber V, Hetz C (2021) Proteostasis deregulation as a driver of C9ORF72 pathogenesis. J Neurochem 159: 941–957 Doi 10.1111/jnc.15529

81 Tsai JC, Miller-Vedam LE, Anand AA, Jaishankar P, Nguyen HC, Renslo AR, Frost A, Walter P (2018) Structure of the nucleotide exchange factor eIF2B reveals mechanism of memory-enhancing molecule. Science 359: Doi 10.1126/science.aaq0939

82 Tshilenge KT, Aguirre CG, Bons J, Gerencser AA, Basisty N, Song S, Rose J, Lopez-Ramirez A, Naphade S, Loureiro A et al (2023) Proteomic Analysis of Huntington’s Disease Medium Spiny Neurons Identifies Alterations in Lipid Droplets. Molecular & cellular proteomics: MCP 22: 100534 Doi 10.1016/j.mcpro.2023.100534

83 Turner MR, Hardiman O, Benatar M, Brooks BR, Chio A, de Carvalho M, Ince PG, Lin C, Miller RG, Mitsumoto H et al (2013) Controversies and priorities in amyotrophic lateral sclerosis. Lancet Neurol 12: 310–322 Doi 10.1016/S1474-4422(13)70036-X

84 Valenzuela V, Becerra D, Astorga JI, Fuentealba M, Diaz G, Bargsted L, Chacon C, Martinez A, Gozalvo R, Jackson Ket al (2025) Artificial enforcement of the unfolded protein response reduces disease features in multiple preclinical models of ALS/FTD. Molecular therapy: the journal of the American Society of Gene Therapy 33: 1226–1245 Doi 10.1016/j.ymthe.2025.01.004

85 Vattem KM, Wek RC (2004) Reinitiation involving upstream ORFs regulates ATF4 mRNA translation in mammalian cells. Proc Natl Acad Sci U S A 101: 11269–11274 Doi 10.1073/pnas.04005411010400541101 [pii]

86 Verdone BM, Cicardi ME, Wen X, Sriramoji S, Russell K, Markandaiah SS, Jensen BK, Krishnamurthy K, Haeusler AR, Pasinelli P et al (2022) A mouse model with widespread expression of the C9orf72-linked glycine-arginine dipeptide displays non-lethal ALS/FTD-like phenotypes. Sci Rep 12: 5644 Doi 10.1038/s41598-022-09593-z

87 Viera Ortiz AP, Cajka G, Olatunji OA, Mikytuck B, Shalem O, Lee EB (2023) Impaired ribosome-associated quality control of C9orf72 arginine-rich dipeptide-repeat proteins. Brain 146: 2897–2912 Doi 10.1093/brain/awac479

88 Wang M, Kaufman RJ (2016) Protein misfolding in the endoplasmic reticulum as a conduit to human disease. Nature 529: 326–335 Doi 10.1038/nature17041

89 Webster CP, Smith EF, Bauer CS, Moller A, Hautbergue GM, Ferraiuolo L, Myszczynska MA, Higginbottom A, Walsh MJ, Whitworth AJet al (2016) The C9orf72 protein interacts with Rab1a and the ULK1 complex to regulate initiation of autophagy. EMBO J 35: 1656–1676 Doi 10.15252/embj.201694401

90 Westergard T, McAvoy K, Russell K, Wen X, Pang Y, Morris B, Pasinelli P, Trotti D, Haeusler A (2019) Repeat-associated non-AUG translation in C9orf72-ALS/FTD is driven by neuronal excitation and stress. EMBO Mol Med 11: Doi 10.15252/emmm.201809423

91 Wong YL, LeBon L, Edalji R, Lim HB, Sun C, Sidrauski C (2018) The small molecule ISRIB rescues the stability and activity of Vanishing White Matter Disease eIF2B mutant complexes. Elife 7: Doi 10.7554/eLife.32733

92 Yang Y, Liu JJ (2022) Structural LTP: Signal transduction, actin cytoskeleton reorganization, and membrane remodeling of dendritic spines. Curr Opin Neurobiol 74: 102534 Doi 10.1016/j.conb.2022.102534

93 Zhang K, Daigle JG, Cunningham KM, Coyne AN, Ruan K, Grima JC, Bowen KE, Wadhwa H, Yang P, Rigo F et al (2018) Stress Granule Assembly Disrupts Nucleocytoplasmic Transport. Cell 173: 958–971 e917 Doi 10.1016/j.cell.2018.03.025

94 Zhang YJ, Gendron TF, Ebbert MTW, O’Raw AD, Yue M, Jansen-West K, Zhang X, Prudencio M, Chew J, Cook CN et al (2018) Poly(GR) impairs protein translation and stress granule dynamics in C9orf72-associated frontotemporal dementia and amyotrophic lateral sclerosis. Nat Med 24: 1136–1142 Doi 10.1038/s41591-018-0071-1

95 Zhang YJ, Jansen-West K, Xu YF, Gendron TF, Bieniek KF, Lin WL, Sasaguri H, Caulfield T, Hubbard J, Daughrity L et al (2014) Aggregation-prone c9FTD/ALS poly(GA) RAN-translated proteins cause neurotoxicity by inducing ER stress. Acta neuropathologica 128: 505–524 Doi 10.1007/s00401-014-1336-5

96 Zyryanova AF, Weis F, Faille A, Alard AA, Crespillo-Casado A, Sekine Y, Harding HP, Allen F, Parts L, Fromont C et al (2018) Binding of ISRIB reveals a regulatory site in the nucleotide exchange factor eIF2B. Science 359: 1533–1536 Doi 10.1126/science.aar5129

